# The impact of mutational processes on structural genomic plasticity in cancer cells

**DOI:** 10.1101/2021.06.03.446999

**Authors:** Tyler Funnell, Ciara H O’Flanagan, Marc J Williams, Andrew McPherson, Steven McKinney, Farhia Kabeer, Hakwoo Lee, Tehmina Masud, Peter Eirew, Damian Yap, Allen W Zhang, Jamie L P Lim, Beixi Wang, Jazmine Brimhall, Justina Biele, Jerome Ting, Yi Fei Liu, Sean Beatty, Daniel Lai, Jenifer Pham, Diljot Grewal, Douglas Abrams, Eliyahu Havasov, Samantha Leung, Viktoria Bojilova, Richard A Moore, Nicole Rusk, Florian Uhlitz, Nicholas Ceglia, Adam C Weiner, J Maxwell Douglas, Dmitriy Zamarin, Britta Weigelt, Sarah H Kim, Arnaud Da Cruz Paula, Jorge S. Reis-Filho, Yangguang Li, Hong Xu, Teresa Ruiz de Algara, So Ra Lee, Viviana Cerda Llanos, IMAXT consortium, Sohrab P. Shah, Samuel Aparicio

## Abstract

Structural genome alterations are determinants of cancer ontogeny and therapeutic response. While bulk genome sequencing has enabled delineation of structural variation (SV) mutational processes which generate patterns of DNA damage, we have little understanding of how these processes lead to cell-to-cell variations which underlie selection and rates of accrual of different genomic lesions. We analysed 309 high grade serous ovarian and triple negative breast cancer genomes to determine their mutational processes, selecting 22 from which we sequenced >22,000 single cell whole genomes across a spectrum of mutational processes. We show that distinct patterns of cell-to-cell variation in aneuploidy, copy number alteration (CNA) and segment length occur in homologous recombination deficiency (HRD) and fold-back inversion (FBI) phenotypes. Widespread aneuploidy through induction of HRD through *BRCA1* and *BRCA2* inactivation was mirrored by continuous whole genome duplication in HRD tumours, contrasted with early ploidy fixation in FBI. FBI tumours exhibited copy number distributions skewed towards gains, widespread clone-specific variation in amplitude of high-level amplifications, often impacting oncogenes, and break-point variability consistent with progressive genomic diversification, which we termed serriform structural variation (SSV). SSVs were consistent with a CNA-based molecular clock reflecting a continual and distributed process across clones within tumours. These observations reveal previously obscured genome plasticity and evolutionary properties with implications for cancer evolution, therapeutic targeting and response.

## INTRODUCTION

Mutational processes in human cancer arising from endogenous DNA repair deficiencies often lead to genomic instability, resulting in specific patterns of base level substitutions^1^, structural variations (SVs)^2^, copy number alterations (CNAs) and formation of micronuclei^3^. Breakage-fusion-bridge and homologous recombination deficiency (HRD), for example, result in structural variation accrual with specific patterns^2,4–6^, measurable using whole genome sequencing (WGS). Computational machine learning approaches can then reconstruct the generative mutational processes that have been active in the history of the tumour. These approaches have transformed our understanding of cancer genome etiology^4,7–10^, and in turn enable improved stratification to determine prognostic and therapeutic significance for patients^5,6,11^.

Mutational processes have typically been derived from bulk sequencing and thus the ‘active’ impacts of genomic instability in the most recent cell divisions of tumour cells have been understudied. Bulk-derived data are unsuited to separate evolutionary vestigial events in the initial clonal expansion, from contemporaneous events that better reflect ongoing mechanisms of cell to cell genomic diversification. Single cell WGS, on the other hand, can readily decompose clonal and cellular genomic events^12–14^, enabling calculation of CNA structural variant accrual rates over thousands of individual cells. Thus, single cell approaches are poised to unveil the nature of how different mutational processes lead to diversifying the genomes of individual cells in human tumours and how, in turn, they lead to clonal expansions driven by specific copy number event types. As high grade serous ovarian cancers (HGSCs) and triple negative breast cancers (TNBCs) are both patterned by profound genomic instability and mutational processes defined by SVs^4,5,9,15,16^, we accordingly set out to investigate how SV-associated mutational processes impact genome replication and cell-to-cell genome variation in these cancers.

## RESULTS

### SV mutational processes are common across HGSC and TNBC

We began by profiling a ‘meta-cohort’ of 309 patients comprising 170 HGSCs and 139 TNBCs with bulk tumour-normal paired WGS to infer the distribution of mutational processes across these cancers (**Fig. 1, Supplementary Table 1**). 106 TNBC and 22 HGSC genomes were newly sequenced for this study, and were combined with previously published HGSC^5,17–19^ and TNBC^12,20–23^ datasets. We applied a previously described correlated topic model machine learning approach (MMCTM)^4^, jointly analysing and modeling statistical associations across 10 single-nucleotide variant (SNV) and 9 SV mutational patterns (**Extended Data Fig. 1, Supplementary Table 1, Supplementary Table 2**). After clustering using the signature probabilities as features, HGSC and TNBC samples were interspersed among 11 subgroups, indicating the presence of shared mutational processes (**Fig. 2a, Extended Data Fig. 2, Supplementary Table 3**). We identified two types of HRD signature containing tumours, enrichment of *BRCA1* mutation and patterns reflecting short/medium tandem duplications (HRD-Dup; cluster 1, n=110), as well as enrichment for *BRCA2* mutation with interstitial deletions (HRD-Del; cluster 3, n=29). APOBEC activity defined a third group (clusters 5, 6; n=20) while loss of *CDK12* and large tandem duplications (TD; cluster 4, n=18), *CCNE1* amplification enriched foldback inversions (FBI; clusters 2, 7, 8, 9; n=63), and large deletions (L-Del; cluster 11, n=38) comprised the remaining tumours as distinct from HRD. These signature types were found in both HGSC and TNBC tumours with varying proportions (**Fig. 2b**) and accumulation of SNVs or SVs (**Fig. 2c**). Prognostic analysis in HGSCs was consistent with previous results^4,5^ (p-value: 0.0021, **Fig. 2d**), with HRD-Del at higher median survival than HRD-Dup, followed by FBI and TD with the worst median survival (**Fig. 2e**, p-value: 0.003). For TNBCs, no significant differences were found between signature types (**Extended Data Fig. 3a,b**). However, a Cox proportional hazards model fit to standardized signature probabilities found that the L-Del (large deletions) signature was associated with an increased overall and progression-free survival hazard ratio (**Extended Data Fig. 3c,d, Extended Data Fig. 4**).

**Figure 1.**
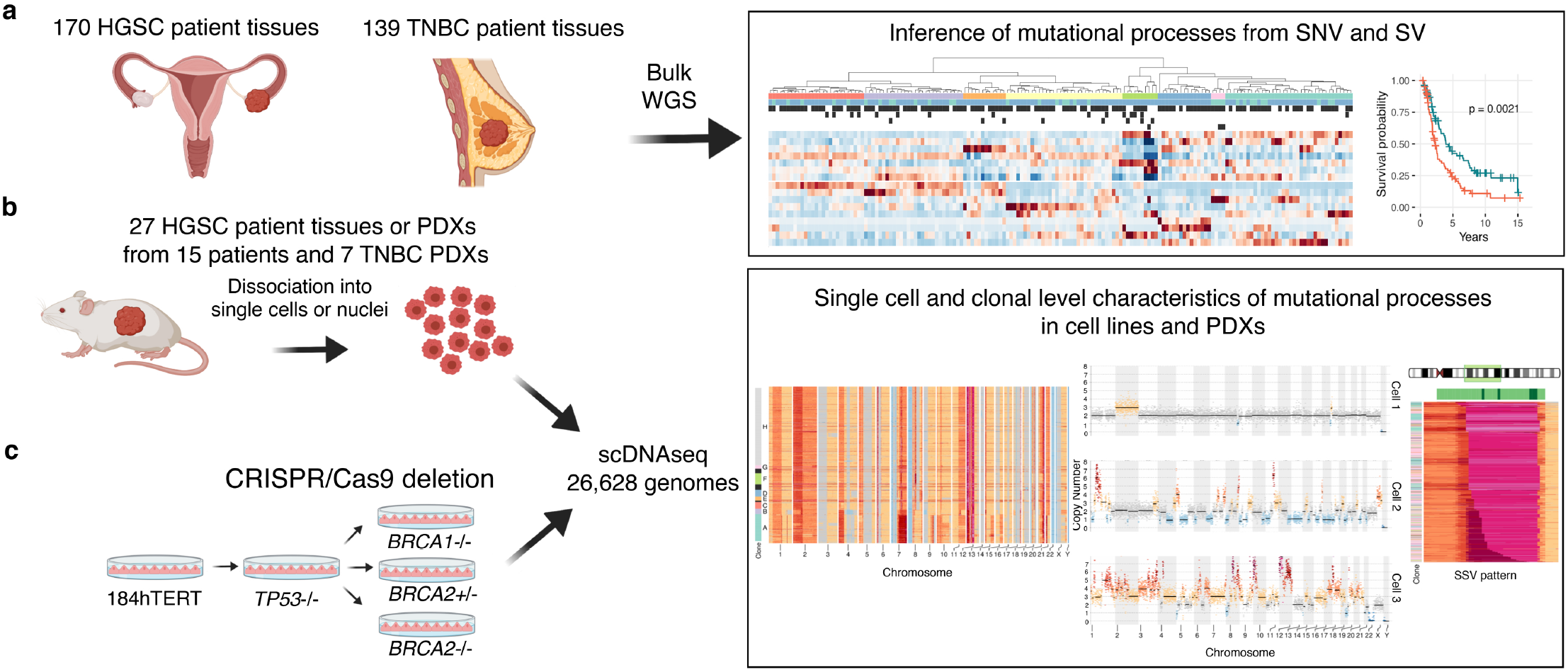
Experimental and cohort design. a) 139 TNBC and 170 HGSC matched normal and tumour bulk whole genomes were grouped according to their SNV and SV mutations using an MMCTM model and prognostic analysis performed on resulting subgroups. b) Single cell genomes from patient or PDX tissue from 22 individual TNBC and HGSC patients from the meta-cohort in a), as well as isogenic 184-hTERT cell lines (WT, or with *TP53, BRCA1* or *BRCA2* mutations) c) were used to examine rates of mutations including chromosomal missegregation, gains, losses, high level amplifications, and serriform structural variants (SSV) at single cell resolution and within clonal populations.

**Figure 2.**
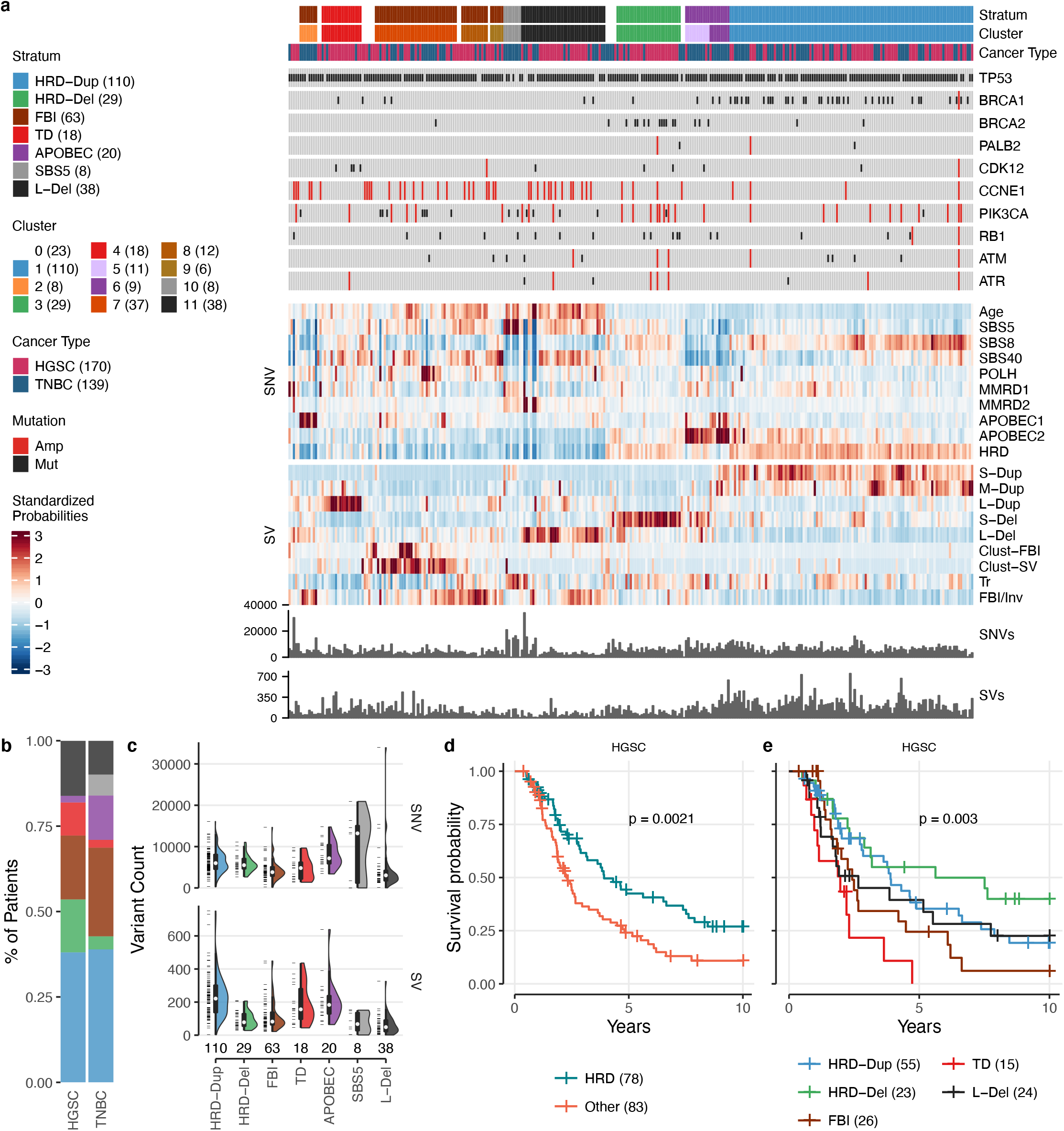
Meta-cohort signature analysis of 139 TNBC and 170 HGSC bulk whole genomes. a) Heatmap representing individual patients as columns, annotation tracks (top) including cancer type and mutation status of key genes, standardized signature probabilities of SNVs and SVs (middle) and event counts (bottom). b) Signature type (see stratum annotation track) proportions by cancer type. c) SNV and SV count distributions per signature type. Boxplots within violins represent the median (white dot), 1^st^ and 3^rd^ quartiles (hinges), and the most extreme data points no farther than 1.5x the interquartile range from the hinge (whiskers). # samples indicated below violins. Data points shown left of violins. Kaplan-Meier survival probability of HGSCs faceted by d) HRD and e) more granular signatures.

### BRCA1 and BRCA2 mutations generate distinct rates of polyploidy and segmental aneuploidy

Structural variation differences between *BRCA1* and *BRCA2* associated tumours observed in bulk prompted us to delineate how HRD-Dup and HRD-Del mutational processes lead to accrual of alterations at the single cell level. We first investigated an isogenic *in vitro* system, through generating loss of function *TP53^21^, TP53/BRCA1* and *TP53/BRCA2* alleles from diploid 184-hTERT mammary epithelial cells^24^ using CRISPR-Cas9 editing (**Supplementary Table 4, Extended Data Fig. 5, Extended Data Fig. 6, Extended Data Fig. 7**). After 20-30 passages, we then generated single cell WGS libraries (DLP+) (median 0.04x coverage, **Extended Data Fig. 8**) from each genotype combination as follows: 184-hTERT^wt^ (wild-type, WT, n=877 genomes), 184-hTERT^TP53-/-^ (TP53^-/-^, n=1722), 184-hTERT^TP53-/-;BRCA1-/-^ (BRCA1^-/-^, n=380), 184-hTERT^TP53-/-;BRCA2-/-^ (BRCA2^-/-^, n=753) and 184-hTERT^TP53-/-;BRCA2+/-^ (*BRCA2*^+/-^, n=605) (**Fig. 3a, Extended Data Fig. 8, Supplementary Table 5, Supplementary Table 6**).

**Figure 3.**
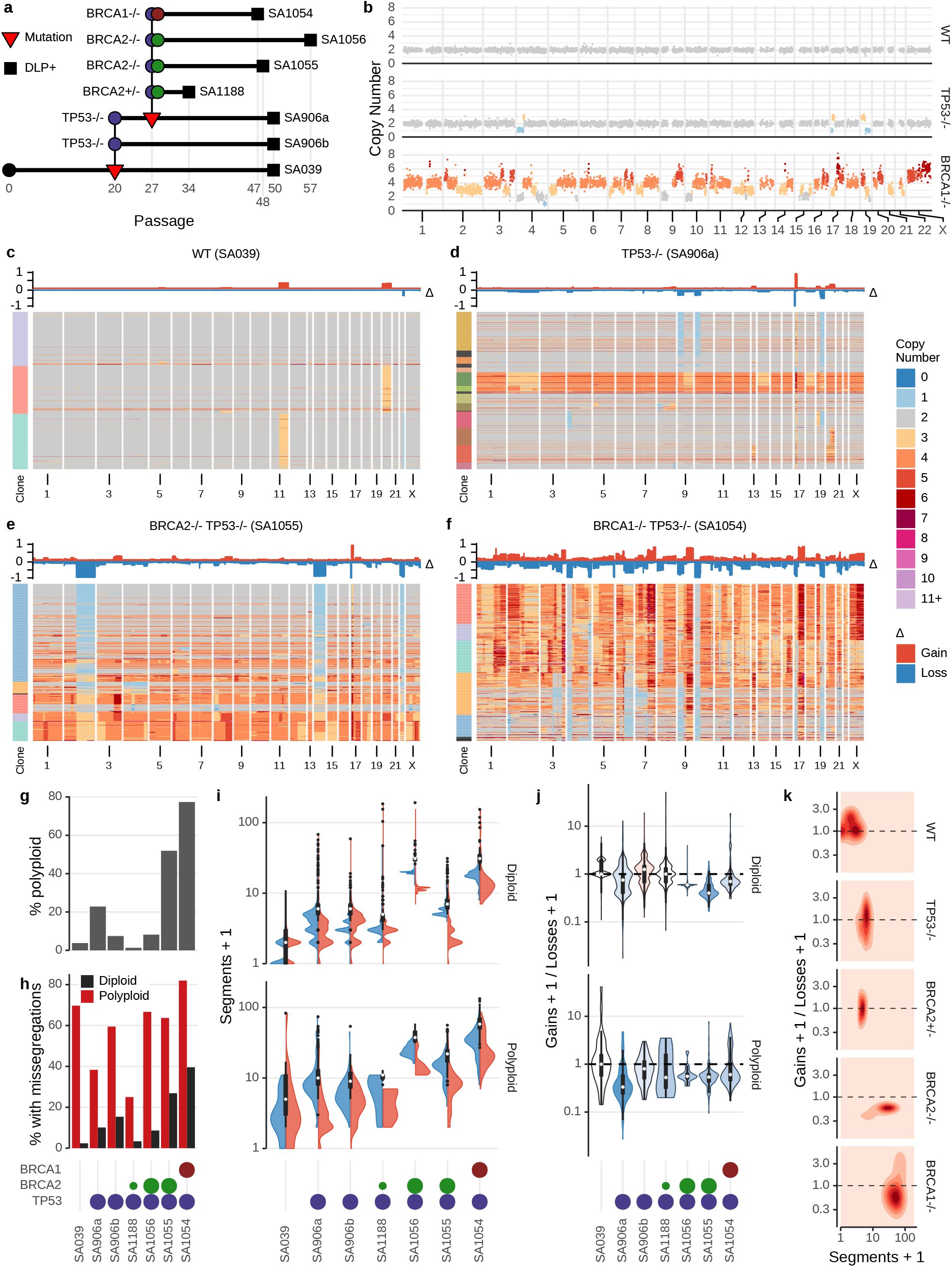
Single cell genome properties of CRISPR/Cas9-derived isogenic genotypes of 184-hTERT mammary epithelial cell lines. a) Genotype lineage diagram showing *WT→TP53→BRCA1/2* interventions. b) Example single cell genomes measured with DLP+. c-f) Heatmap representations of copy number profiles of cell populations of WT, *TP53*^-/-^, *BRCA2*^-/-^ and *BRCA1*^-/-^ lineages. Comparisons of g) rate of polyploidization h) rate of missegregation i) distribution over number of segments with gains (red) and loss (blue) j) distribution over ratio of gains / losses and k) 2-dimensional distribution of gain/loss ratio as a function of number of segments. All boxplots within violins represent the median (white dot), 1^st^ and 3^rd^ quartiles (hinges), and the most extreme data points no farther than 1.5x the interquartile range from the hinge (whiskers).

Per-cell copy number profiles (*e.g*. **Fig. 3b**) exhibited a progressive increase in CNA rates as a function of *TP53* and *BRCA1/2* loss. The majority of WT cells were diploid, with increasing aneuploidy of *TP53*^-/-^, *BRCA2*^-/-^ and *BRCA1*^-/-^ respectively. In *BRCA1*^-/-^, the majority of cells had also undergone a whole genome duplication (**Fig. 3c-f, Supplementary Table 7**). Increasing aneuploidy rate was accompanied by increase in rates of polyploidization (**Fig. 3g**) and chromosomal missegregation events (**Fig. 3h, Supplementary Table 8**). Notably, overall missegregation rates were higher in polyploid cells than diploid cells (p-value: 0.00066), likely reflecting increased tolerance to chromosomal alteration on a whole genome doubled background.

We next computed median per-cell segmental alteration counts (**Fig. 3i, Supplementary Table 9**) and found *BRCA1*^-/-^ genomes (53 events per cell) harboured higher rates of segmental alteration than *BRCA2*^-/-^ (30 and 10), *BRCA2*^+/-^ (4) *TP53*^-/-^ (5) or WT cell lines (1) (all p-values: <10e-10). As for missegregations, polyploid cells harbored a greater variance in segmental alterations (median 9.3X increase), again reflecting increased tolerance for genomic plasticity after whole genome duplication. We then compared the ratio of gains to losses, assuming unbalanced ratios would indicate tolerance away from neutrality. The ratio was balanced in the WT cells and the *BRCA2*^+/-^, however *BRCA1*^-/-^ and *BRCA2*^-/-^ cells exhibited skewed ratios towards losses in diploid cells (p-values: <0.05) (**Fig. 3j**) and all mutant libraries showed skewing towards losses in polyploid cells. The gain/loss ratio plotted as a function of segmental aneuploidies revealed both an increase of aneuploidy and a higher relative shift towards losses for *BRCA1*^-/-^ compared to *BRCA2*^-/-^ (**Fig. 3k**). We suggest this may reflect distinct evolutionary patterns in *BRCA1* and *BRCA2* deficient cells such that despite skewing towards accumulation of losses at the single cell level, human cancers nevertheless exhibit tandem duplication enrichment at the bulk genome level in *BRCA1* associated cancers, but enrichment of interstitial deletions in *BRCA2* associated cancers^4,8^ (see **Fig. 2a**). Thus, cell-level genomic consequences of *BRCA1* and *BRCA2* inactivation induced higher whole genome doubling, higher missegregation and higher rates of segmental aneuploidies relative to the *TP53*^-/-^ background by orders of magnitude, indicating BRCA loss as a major diversifying process. Higher background rates of genomic instability with *BRCA1* compared to *BRCA2* loss may further explain differences in cisplatin response. Our results here comparing *BRCA2*^+/-^ and *BRCA2*^-/-^ reinforce that bi-allelic loss is required to impact rates of genomic instability.

### Distinguishing HRD and FBI single cell genomes in tumours

Having contrasted *BRCA1* and *BRCA2* signatures *in vitro*, we next investigated how structural variation processes were reflected in single cell genomes of human tumours. We selected 22 cases (15 HGSC and 7 TNBC) whose primary tumours were classified into signature types in the meta-cohort (**Fig. 2a, Supplementary Table 6**) and generated patient derived xenografts (PDXs) from 18 of these tumours over a multi-year period using subcutaneous engraftment (**Supplemental methods**). DLP+ libraries from single nuclei or cell suspensions of HRD-Dup (n=8), FBI (n=9), TD (n=2), L-Del (n=2), and APOBEC (n=1) PDX and patient tissues yielded a total of 22,291 genomes (median 458 per series), and median 1.69 million reads per genome (median 0.05X coverage, IQR 0.04, **Extended Data Fig. 8, Supplementary Table 5, Supplementary Table 6**). Pseudobulk analysis, which merges single cell whole genome data into a simulated bulk profile^12^, yielded CNAs and mutations consistent with matched bulk WGS from the primary tumours (**Extended Data Fig. 9**). Single cell copy number profiles (**Fig. 4a**) and population level analysis (**Fig. 4b,c**) indicated broadly distinct patterns of genomic alteration between HRD-Dup and FBI tumours. All tumours harbored single cell level events, consistent with a background process accruing segmental alterations. Notably, FBI tumours were either majority diploid (n=3), or majority polyploid (n=6), consistent with a fixed baseline ploidy on initial clonal expansion without ongoing whole genome duplication (**Supplementary Table 7**). In contrast, HRD-Dup tumours exhibited mixed ploidy distributed over clones (2.5X higher mean Shannon entropy *vs*. FBI, p-value: 0.011, **Fig. 4d, Extended Data Fig. 10**) suggestive of ongoing whole genome duplication without fixation from a baseline diploid state.

**Figure 4.**
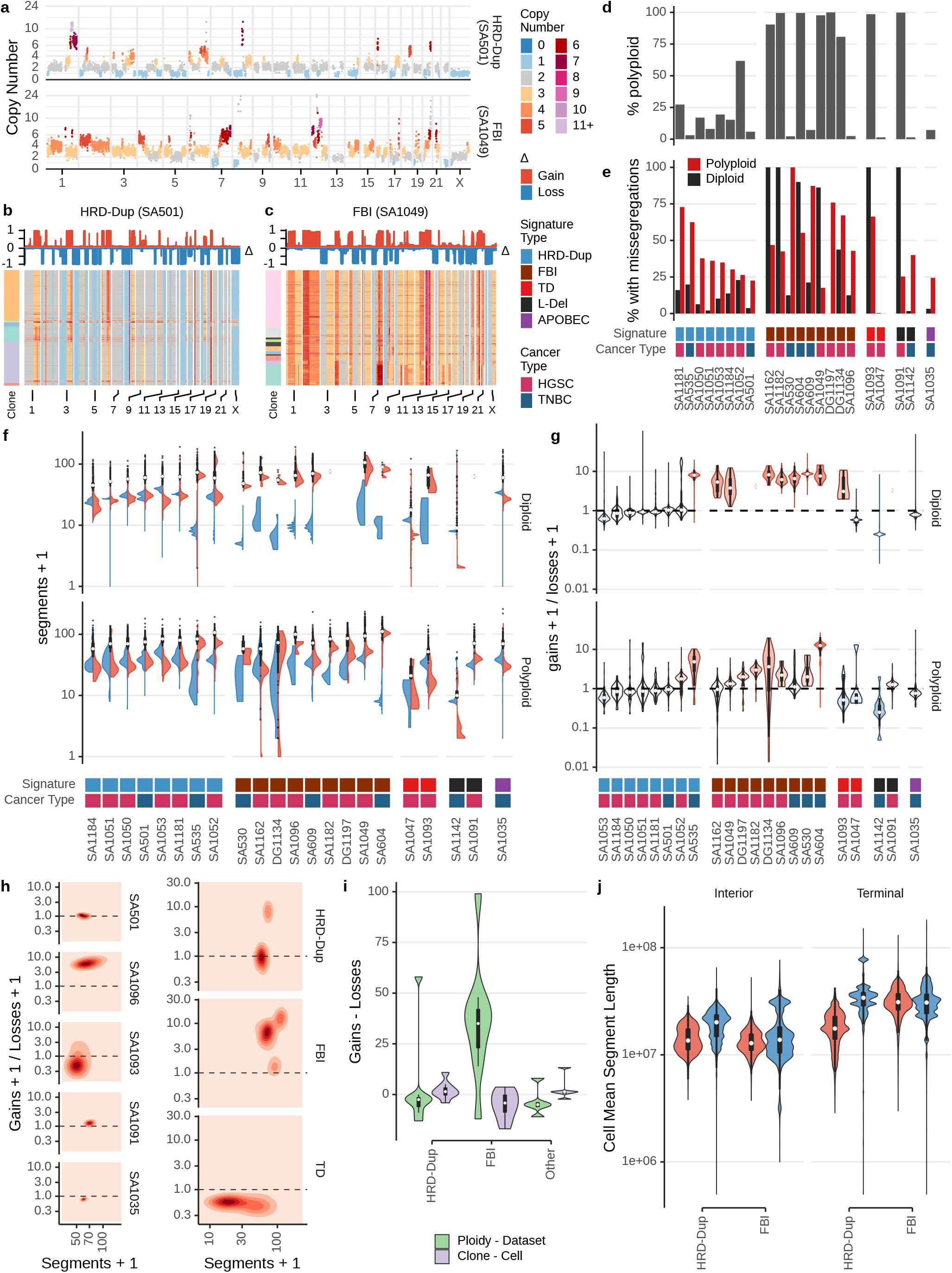
Comparison of rates of accumulation of cell specific events in FBI *vs*. HRD-Dup in PDX models and patient tissues. Segment plot a) and heatmaps b-c) showing chromosomal copy number of single cells from TNBC (SA501) and HGSC (SA1049). d-e) Proportion of cells with missegregation events with respect to ploidy state. (f-g) Chromosomal gains and losses across different ploidy states and mutational signature grouping. h) Relationship between gain/loss ratios and number of gained or lost segments for representative datasets from each signature type (left) and all HRD-Dup, FBI, or TD datasets (right). i) Difference in chromosomal gains and losses for copy number profiles at the dataset level relative to ploidy (green) and cell level relative to clone copy number profiles (purple). j) Mean gain (red) and loss (blue) segment lengths of interior and terminal chromosome regions in HRD-Dup and FBI subgroups. All boxplots within violins represent the median (white dot), 1^st^ and 3^rd^ quartiles (hinges), and the most extreme data points no farther than 1.5x the interquartile range from the hinge (whiskers).

Diploid FBI cells exhibited higher overall rates of missegregation relative to HRD-Dup (**Fig. 4e**, p-value: 0.066, **Supplementary Table 8**). Patterns of changes in the gain/loss ratio as a function of number of segments indicated FBI tumours accrue gains at a significantly higher rate than HRD-Dup (**Fig. 4f,g**; p-value: <10e-10), with orders of magnitude more skewing of the gain/loss ratio (**Fig. 4h**). The increase of segmental gains in FBI tumours was most pronounced when relative to baseline ploidy of the tumours (**Fig. 4i**, p-value: 0.0093, **Supplementary Table 10, Supplementary Table 11**). The changes between clone consensus profiles and single cell profiles represent more recent events, and were generally more balanced in both HRD-Dup and FBI cases. Notably, FBI segment lengths at chromosomal termini were 1.8x longer in FBI, while HRD-Dup exhibited longer segments internal to the chromosome (**Fig. 4j**, p-value: <10e-10), consistent with different originating sites of copy number events; this may reflect differential telomere dysfunction across the two mutational processes. Thus, considering whole genome duplication, overall rates of segmental aneuploidy, gain/loss ratio, and positional locations of segments along the chromosome, FBI and HRD-Dup tumours showed strikingly different CNA accrual patterns at single cell level.

### Copy number amplitude and serriform breakpoint variation over single cells is common in FBI tumours

As FBI tumours accrued copy number gains at orders of magnitude higher rates than HRD cancers, we next studied how these gains led to regions of high-level amplification (HLAMP, here defined as >10 copies). HLAMPs are a major source of oncogenic potential and co-localised FBIs with HLAMPs are prognostically significant in HGSC^5^. To determine if HLAMPs are fixed, or dynamic in the evolution of these cancers, we enumerated all HLAMPs present in at least 10 cells in each dataset and measured cell-to-cell variation in amplitude. We found that variation in the amplitude across clonal populations within tumours was common (example events shown in specific clones of SA1049, an FBI tumour, at the *KRAS* locus (**Fig. 5a**) and genome wide (**Fig. 5b**)), revealing new complexity in the cell-to-cell composition of HLAMPs that would be obscured by bulk analysis. Comparisons by mutational process showed FBI tumours with 3.4x higher median copy variance than the other tumours (**Fig. 5c-g**; p-value: 0.00054, **Supplementary Table 12**), consistent with continual plasticity of the amplitude of HLAMPs as a general property of FBI. Furthermore, we noted amplitude variation in HLAMPs impacting known oncogenes including *ERBB2* (DG1197), *KIT* (DG1197), *KRAS* (SA1049, SA604), *MYC* (SA1184, SA1035, SA1051), *CCNE1* (DG1134, SA1162, SA604), and *FGFR1* (SA1049, SA535), suggesting that clone-specific variations in amplitude would likely yield clone-specific phenotypic variation (**Fig. 5d, Supplementary Table 13**).

**Figure 5.**
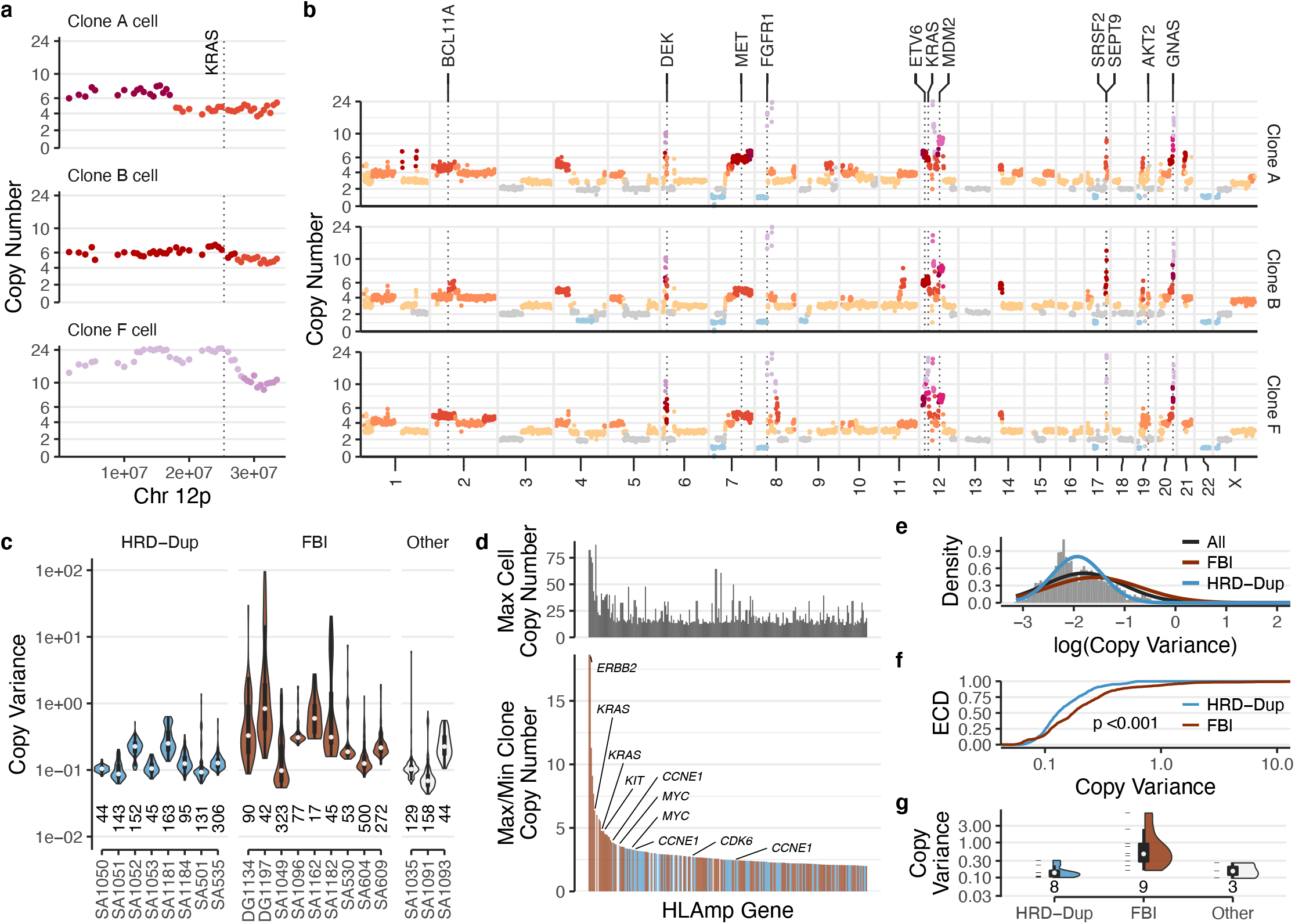
HLAMP copy number variance. a) Chromosome 12p copy number for 3 cells from SA1049 clones A, B, and F, with *KRAS* gene locus indicated with dotted line. b) Clone whole genome consensus copy number profiles for SA1049 clones A, B, and F. Selected genes overlapping HLAMP regions indicated with dotted lines. c) Copy number variance across cells for HLAMP bins within each dataset. Datasets without HLAMPs not shown. d) *bottom*: Clone max/min copy number ratio of cancer genes overlapping HLAMP regions. Genes across all cancer datasets with ratio > 2 shown. Colours as per c). *Top*: Max cell copy number for genes below. e) Copy variances for HLAMPs in all cancer datasets. Normal density shown in *black*. Normal density fit to HRD-Dup and FBI HLAMP copy variances: *blue* and *brown*, respectively. X-axis upper limit truncated to 2. f) Empirical cumulative densities of HRD-Dup, *blue*, and FBI, *brown*, HLAMP copy variances. P-value for KS test indicated. X-axis upper limit truncated to 10. g) Mean copy number variances for HLAMPs in each dataset, grouped by signature type. All boxplots within violins represent the median (white dot), 1^st^ and 3^rd^ quartiles (hinges), and the most extreme data points no farther than 1.5x the interquartile range from the hinge (whiskers). # data points indicated below violins. Data points shown left of violins.

In addition to variation in amplitude of HLAMPs, we observed extensive within-tumour cell-to-cell variation in the genomic locations of breakpoints of CNA events. The precise boundaries of CNAs from cell-to-cell yielded a pattern which we termed “serriform structural variants” (SSV) (**Fig. 6a**; heatmaps 3, 4, 5 from left, **Supplementary Table 14**). This pattern typically consisted of a modal breakpoint across cells, with ‘tails’ reflecting either a progressive accumulation or ‘erosion’ away from the modal breakpoint. The aggregate, consensus copy number profiles over cells across the complete SSV regions (analogous to what would be seen in bulk sequencing libraries), revealed sloping copy number changes between integer values indicative of an averaged signal with underlying variance. We noted that SSVs were not a ubiquitous signal (**Fig. 6a**; heatmaps 1, 2 from left) such that many events had conserved breakpoints across cells. Moreover, visualization of raw GC-corrected read counts further supported the serriform pattern, ruling out analysis artefacts (**Fig. 6a**, bottom grayscale heatmaps). We computed serration scores to identify the relative degree of variation in breakpoints across cells in each cancer with a generalized linear model accounting for covariate features including: the number of cells with the event, the copy number of adjacent segments, and the length of the higher copy number segment (**Extended Data Fig. 11**, see **Methods**). Comparison of distributions of serration indicated FBIs with high degrees of breakpoint variance in SSV regions relative to HRD-Dup (**Fig. 6b,c**, p-value: 1.1e-09). Moreover, using a threshold of the overall distribution (z-score 1.5) enumerated more highly variable SSV regions in FBI tumours (**Fig. 6d,e**; median 4 *vs*. 0, p-value: 0.057, 49 total).

**Figure 6.**
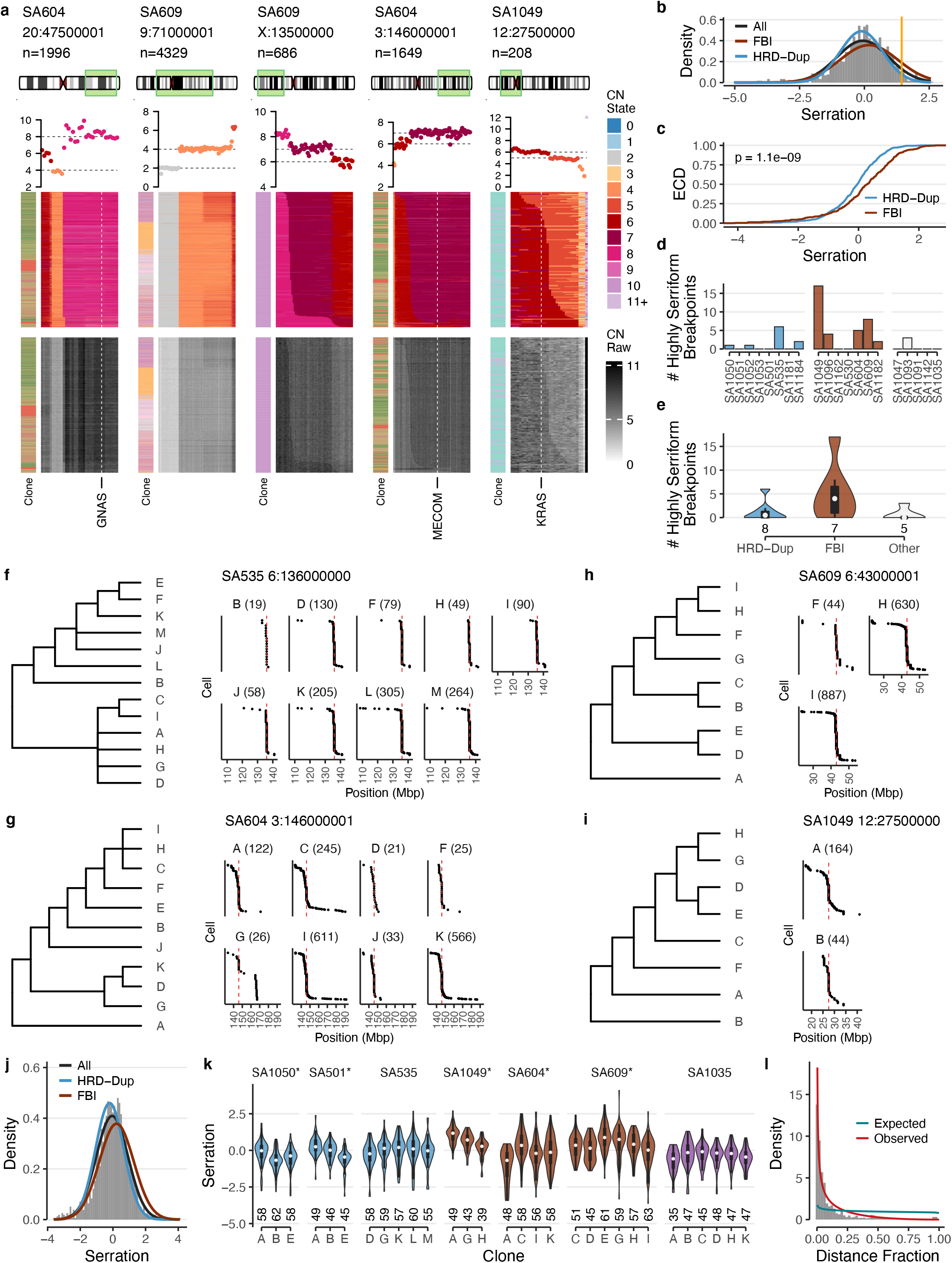
Breakpoint serriform variability. a) Copy number heatmaps showing variation in breakpoint location across cells. *Top to bottom*: dataset, breakpoint location, and # cells; ideogram indicating chromosome region shown in heatmap; average copy number across cells in heatmap, with breakpoint-adjacent segment copy number states indicated with *dotted black* lines; copy number states inferred from HMMCopy; GC-corrected read counts. Heatmap x-axis: genomic bins; y-axis: cells with the indicated breakpoint. Selected gene locations indicated with *white* dotted lines. b) Breakpoint serration distribution for all cancer datasets. Normal density shown in *black*. Normal density fit to HRD-Dup and FBI breakpoint serration scores: *blue* and *brown*, respectively. c) Empirical cumulative densities of HRD-Dup, *blue*, and FBI, *brown*, breakpoint serration. P-value for KS test indicated. d) # highly serriform breakpoints per dataset. DG1134, DG1197 not shown (see Methods) e) # highly serriform breakpoints per dataset, grouped by signature type. f-i) Clone phylogenies and breakpoint locations for example events in SA535, SA604, SA609, and SA1049. Dataset and example breakpoint event coordinates are shown above the breakpoint location plots. Event location indicated with dashed vertical line. Breakpoint locations are split by cell into their corresponding clones, with cell count noted in parentheses. j) Serration for each breakpoint in each tumour clone across all cancer datasets. Densities for all breakpoints, and those in HRD-Dup or FBI tumours are shown with black, blue and brown lines. HRD-Dup vs. FBI KS-test p-value: <10e-10. k) Clone-level breakpoint serration for datasets with ≥3 clones with ≥3 breakpoints. Cancers with significant differences (ANOVA p-value <0.05) in serration between clones are indicated with an asterisk next to the dataset label. l) Per-cell breakpoint distances from modal positions (>5% of cells), as a fraction of the longest distance per event for all highly serrated breakpoints from d). Beta distribution fits for the observed (red, α=0.45, β=3.10) and expected (blue, α=0.89, β=1.00) distance distributions. All boxplots within violins represent the median (white dot), 1^st^ and 3^rd^ quartiles (hinges), and the most extreme data points no farther than 1.5x the interquartile range from the hinge (whiskers). # data points indicated below violins.

We next examined whether serriform patterns were specific to genomically defined clones within tumours or more distributed as indicators of a selected or distributed process. We identified cell clusters as first approximations to clones, computed clone phylogenies, and examined serriform regions at a clone level. For clonal events, we noted that the serriform pattern was present in all major clades of the phylogeny (**Fig. 6f,g**). For clade-specific events, the serriform pattern was present in all clones in the clade of interest (**Fig. 6h,i**). In general, distributions of serration taken over clones were again higher in FBI than HRD-Dup (**Fig. 6j**). Comparison of score distributions within tumours indicated moderate variation from clone to clone, however the serriform patterns were present in all clones across the phylogenies (**Fig. 6k**). The presence across clones of the serriform pattern is consistent with a parallel and distributed process which continues after evolutionary branching and selection. Rare breakpoint positions (<5% of cells) cluster around modal breakpoints, with rare position density decaying with increasing distance from modal breakpoints (**Fig. 6l**). This suggests the serriform process is analogous to a background point mutation rate and could be leveraged as a molecular clock-like process.

Together, these results demonstrate that the underlying mutational process creates increased genomic plasticity both in amplitude of HLAMPs and in cell-specific breakpoints, which reflect a distributed and continual diversification at the copy number level. We expect that FBI tumours therefore create an evolving and broader genomic substrate upon which selection can act in these cancers.

## DISCUSSION

We revealed how structural mutational processes induce cell-to-cell copy number variation in cancer genomes, identifying stark differences between *BRCA1* and *BRCA2* related processes *in vitro*, and between HRD-Dup and FBI human tumours in HGSC and TNBC PDXs and patient tissues. Our findings from 26,628 single cell genomes reshape our understanding of how endogenous mutational processes alter genome architecture in human cancers. Critically our results outline how higher instability in FBI cases manifests in a new understanding of the cellular composition of high level amplifications impacting oncogenes, revealing wide variation in clone specific amplitude. This has direct clinical relevance as some genes including *CCNE1, KIT* and *ERBB2* are targets of therapeutics at various stages of development, and will thus likely lead to incomplete response and/or intrinsic resistance from pre-existing clones^25,26^. Moreover, our results predict clone specific *KRAS* and *MYC* amplifications would impact phenotypic differences within cancers.

We suggest the origins of serriform patterns of copy number breakpoint variation are likely linked to chromosome bridge formation during cell division^27^. Here we illustrate the process is likely distributed across clones in human cancers. This suggests the serriform pattern could be exploited as a molecular clock in FBI tumours, analogous to clock-like processes previously described for point mutations^28–30^. Given the low point mutation rate demonstrated in FBI tumours, the serriform process may therefore be a key determinant to measure etiology and evolutionary timing of oncogenic structural variations such as high level amplifications.

Our results show that both FBI and HRD cancer cells were affected by substantial ongoing genomic instability, yet differed considerably in the nature of variation observed. In HRD-Dup cancers, we found that cell gain and loss distributions were relatively balanced compared to FBI cells which had a clear enrichment of amplifications. In general, FBI cells had greater levels of structural instability, including variation in the amplitudes of HLAMP regions, and in cell-to-cell CNA breakpoint locations. The SSV pattern has been observed *in vitro^27^*, and we have described them in TNBCs and HGSCs for the first time. The relative specificity of SSVs in FBI tumours suggest they could be exploited to refine or develop biomarkers to stratify these high-risk patients. Moreover, the advanced age of diagnosis of FBI patients implicates age-related chromosomal dysfunction as a potential biological origin for SSV genome alteration. For example, whether shortened telomeres initiate the SSV pattern through breakage fusion bridge processes remains a key unresolved question.

In summary, our results and methods open new routes to studying genomic instability in cancer to link DNA damage processes to evolutionary selection, patient stratification and therapeutic targeting.

## METHODS

All methods are described in the Supplemental Materials

## Data availability

Data will be made available for controlled access at EGA upon publication.

## Materials and Correspondence

Requests for materials and correspondence should be addressed to

## Extended Data Figures

**Extended Data Fig. 1.**
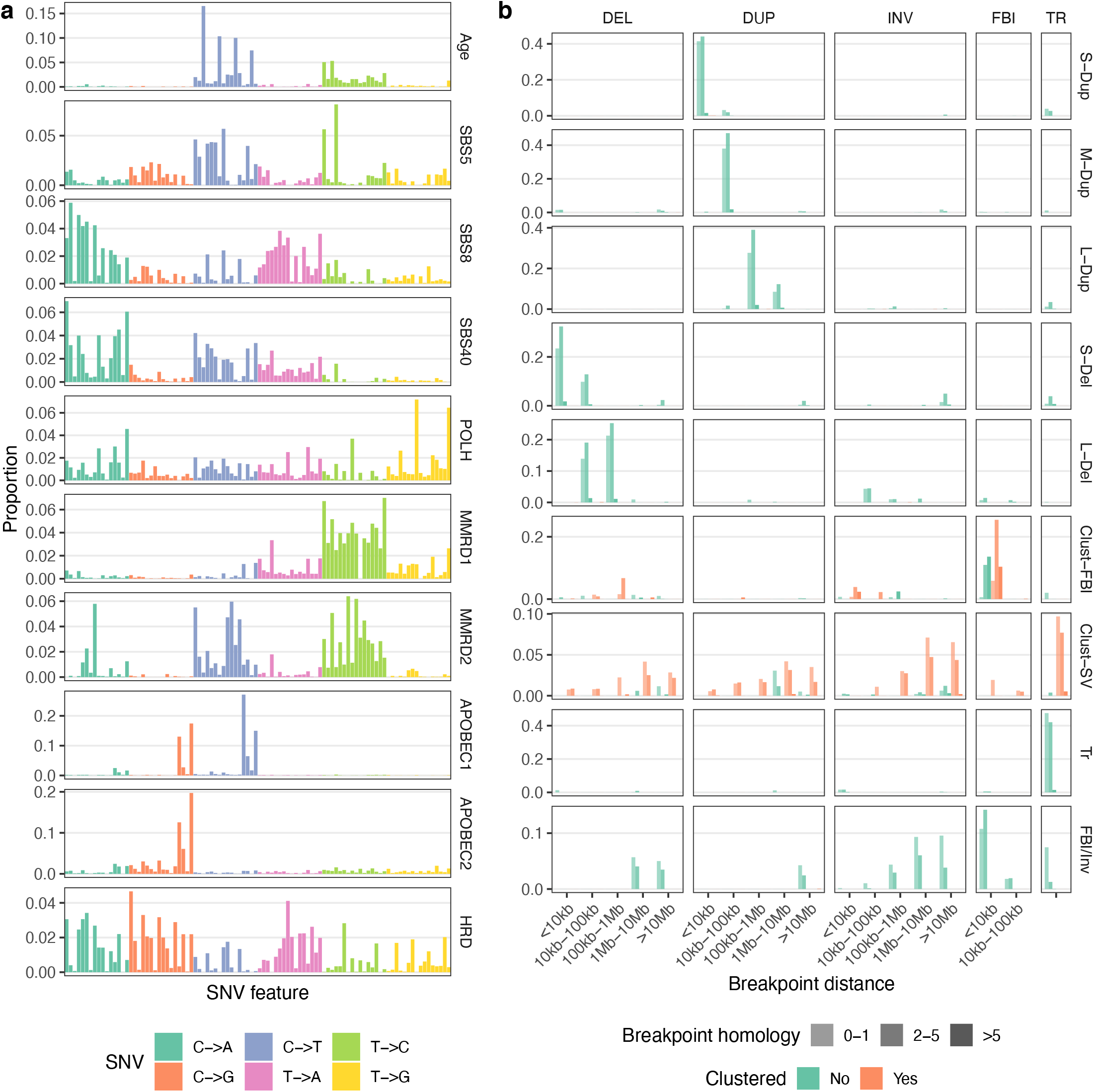
a) SNV and b) SV mutation signatures estimated from HGSC and TNBC bulk tumour mutation catalogues using the MMCTM method. The x-axis in a) is the 96-channel (*i.e*. A[C>A]A, …, T[T>G]T) SNV types.

**Extended Data Fig. 2.**
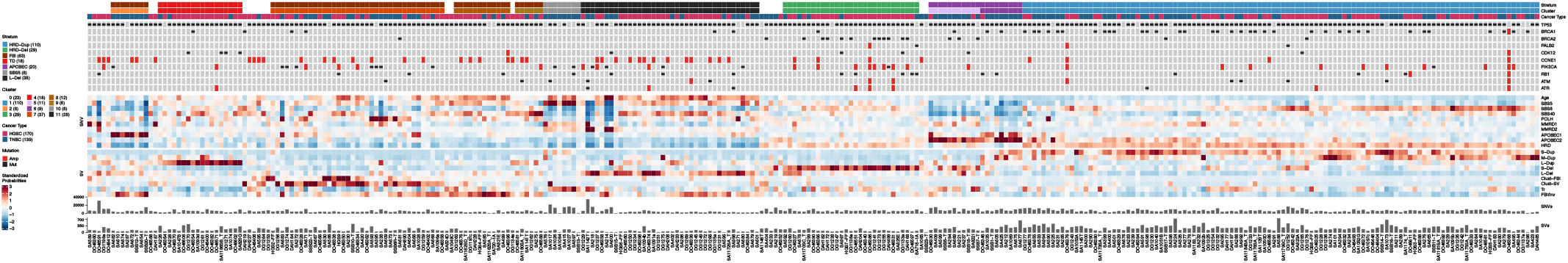
Expanded heatmap of standardized per-sample mutation signature probabilities, with sample identifiers are indicated along the x-axis. Individual patients as columns, annotation tracks (top) including cancer type and mutation status of key genes, standardized signature probabilities of SNVs and SVs (middle) and event counts (bottom).

**Extended Data Fig. 3.**
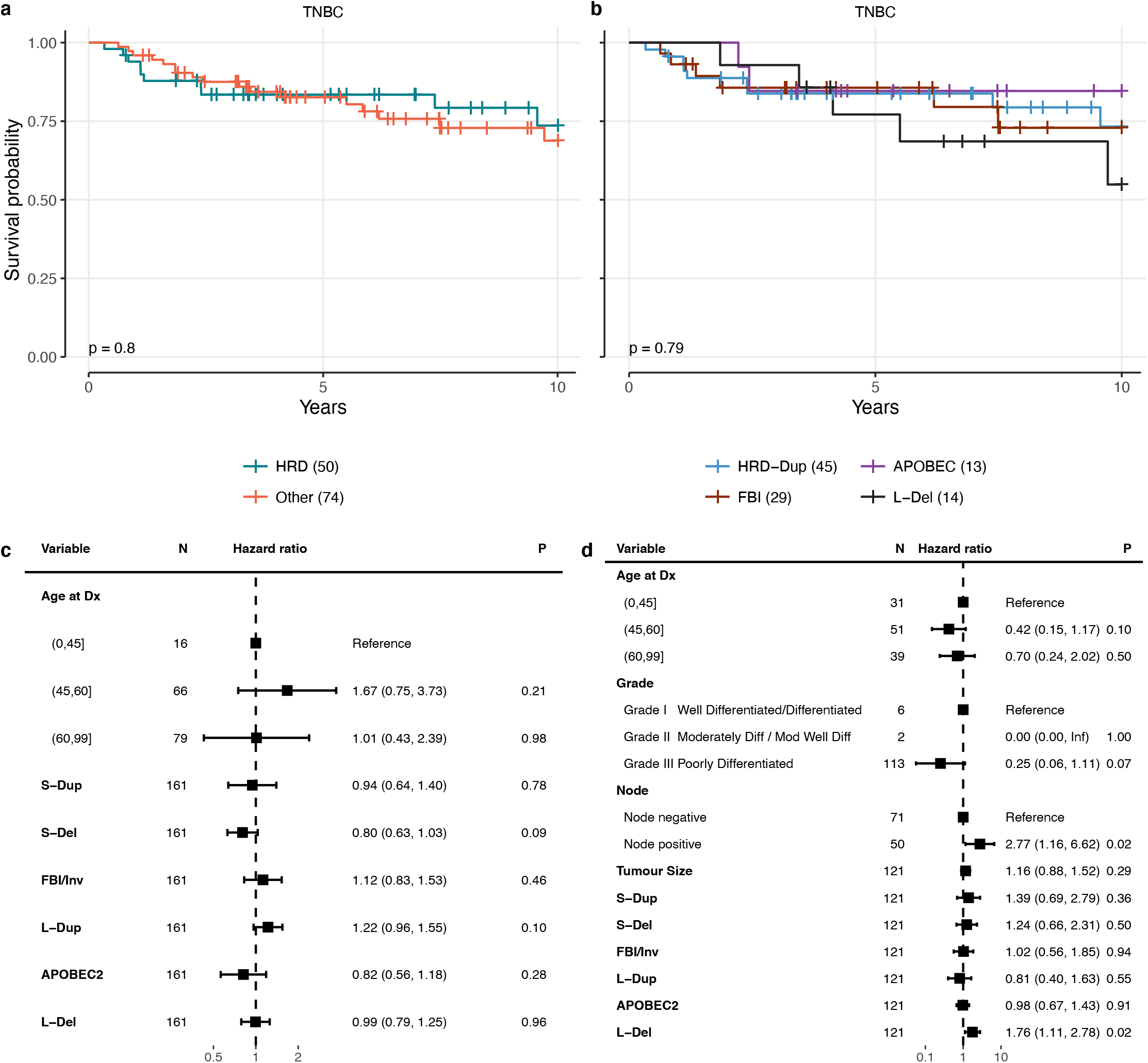
Kaplan-Meier overall survival probability of TNBC faceted by a) HRD and b) more granular signatures. Survival times censored to 10 years. Log-rank test p-value shown in bottom left. Number of cases per group indicated in parentheses. Overall survival Cox proportional hazards models fit with a) HGSC and b) TNBC patient data. Error bars are 95% confidence intervals.

**Extended Data Fig. 4.**
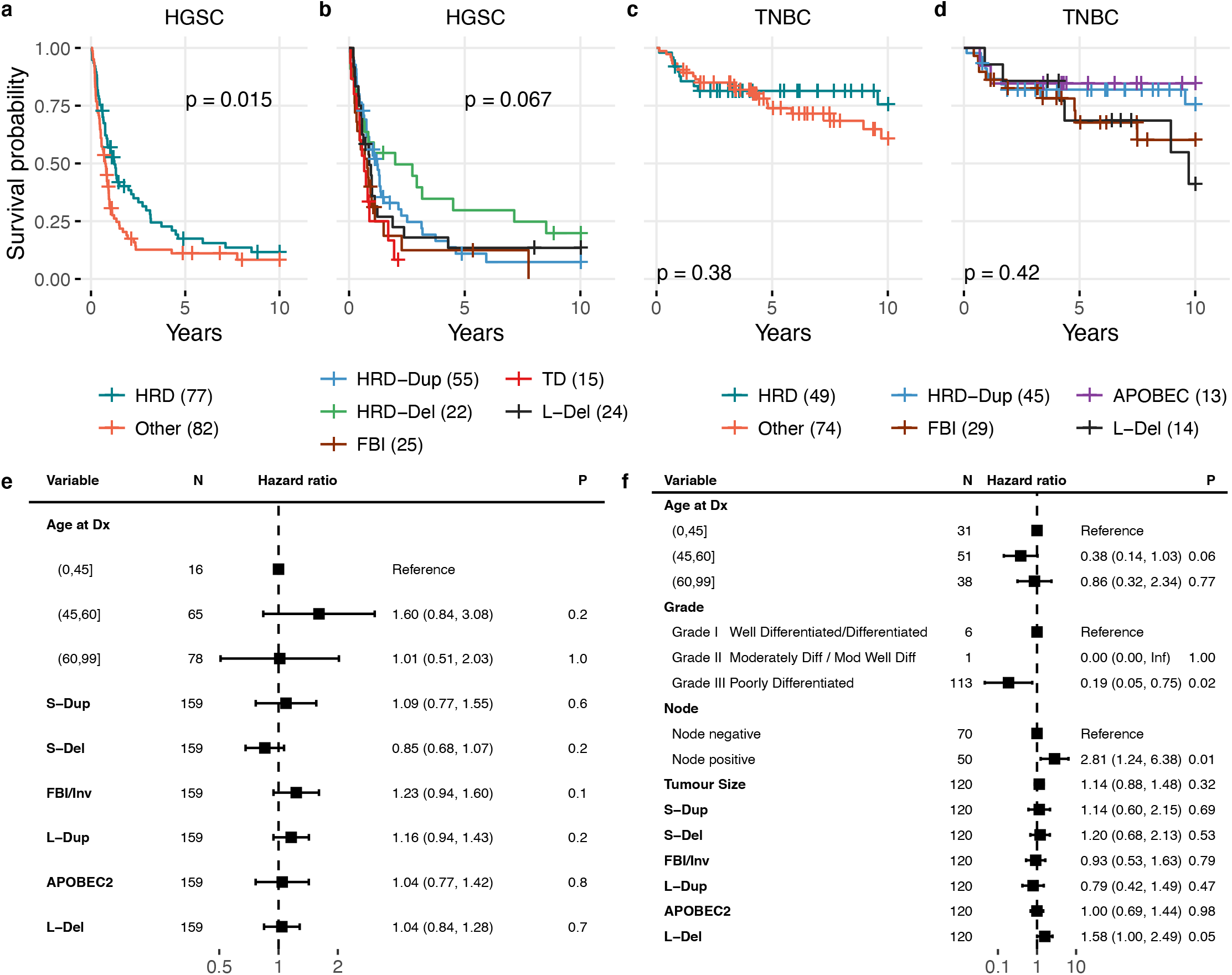
Progression free survival. Kaplan-Meier survival probability of HGSC faceted by a) HRD and b) more granular signatures. Kaplan-Meier survival probability of TNBC faceted by c) HRD and d) more granular signatures. Survival times censored to 10 years. Log-rank test p-value shown. Number of cases per group indicated in parentheses. Cox proportional hazards models fit to e) HGSC and f) TNBC patient progression free survival. Error bars are 95% confidence intervals.

**Extended Data Fig. 5.**
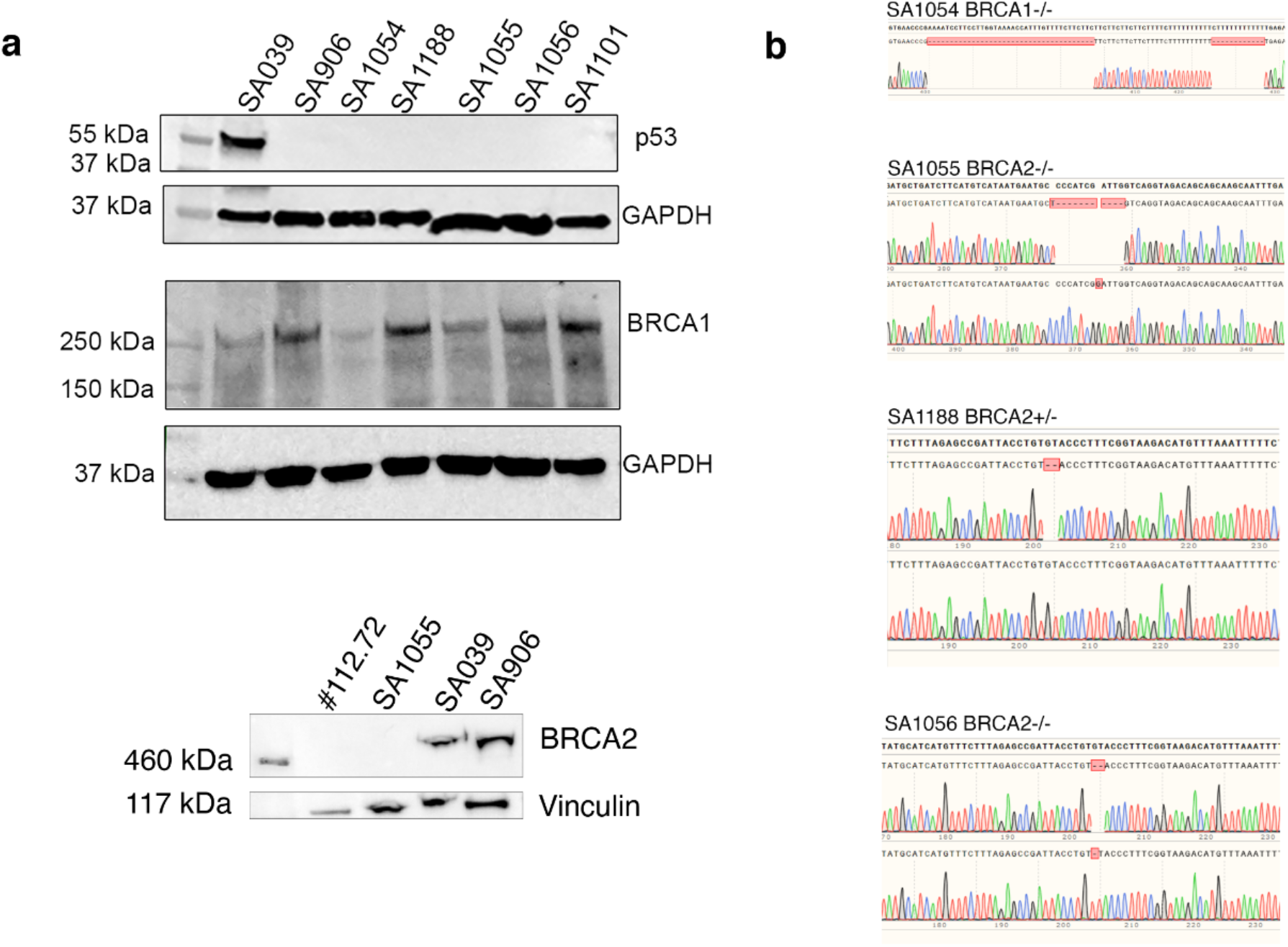
Verification of CRISPR-Cas9 induced genotypes of 184-hTERT cell lines by western blotting for p53, BRCA1 and BRCA2 a), including an additional *BRCA2*-/- clone 112.72. b) Sanger sequencing of TOPO cloned *BRCA1* and *BRCA2* regions.

**Extended Data Fig. 6.**
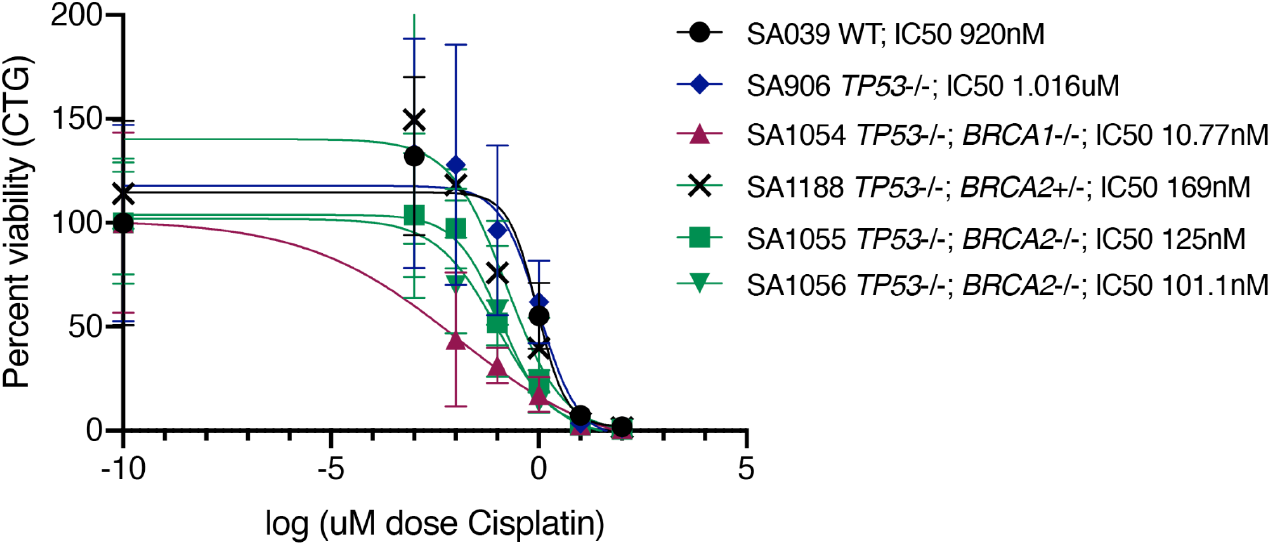
Cell titre glo (CTG) viability assay of cisplatin sensitivity of 184-hTERT cell lines with *TP53*-/-, *BRCA1*-/-, *BRCA2*+/- or *BRCA2*-/- genotypes. Homozygous knockout of *BRCA1* or *BRCA2* as well as heterozygous knockout of *BRCA2* resulted in increased sensitivity of *TP53*-/- 184-hTERT cells to treatment with cisplatin. IC50 determined by nonlinear regression, error bars are standard deviation of 5 replicate wells.

**Extended Data Fig. 7.**
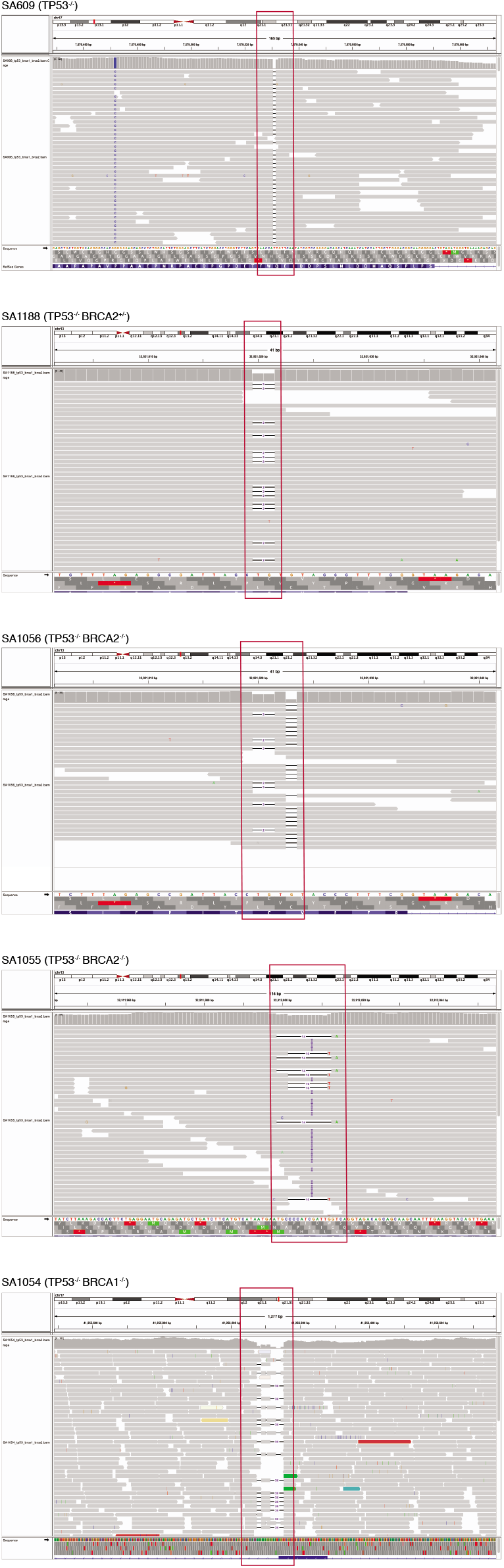
IGV plots of aligned reads from DLP+ data from 184-hTERT cell lines showing the induced mutation in *TP53* for SA906; *BRCA2* for SA1188, SA1056, SA1055; and *BRCA1* for SA1054. Cell lines indicated above each plot. Mutation loci are indicated with red boxes.

**Extended Data Fig. 8.**
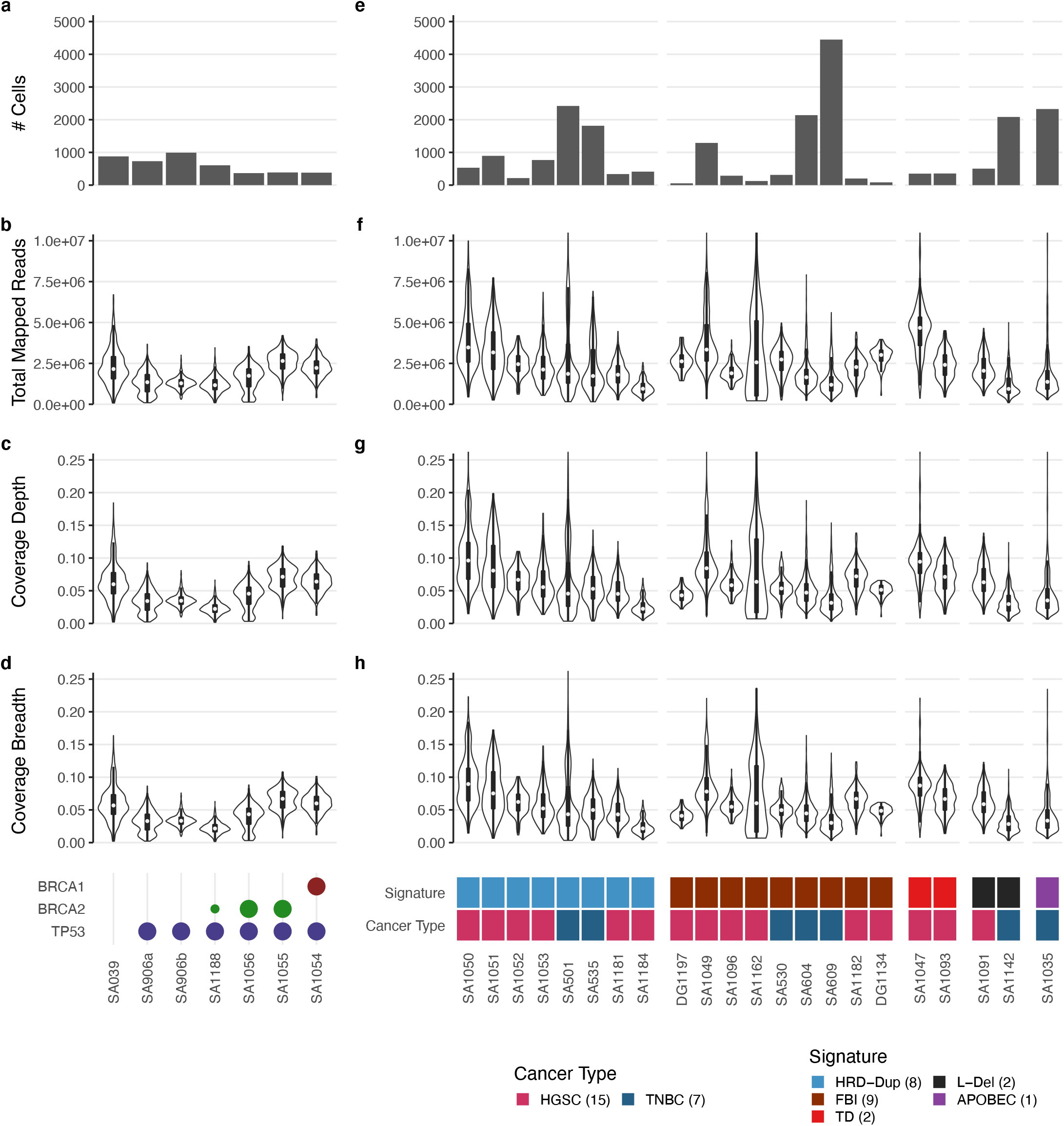
Summary of DLP+ sequencing statistics of data for 184-hTERT cell lines a-d) and HGSC and TNBC tumours e-h). Shared y-axis labels shown at left. The legend for e-h) indicates the number of samples for each cancer and signature type.

**Extended Data Fig. 9.**
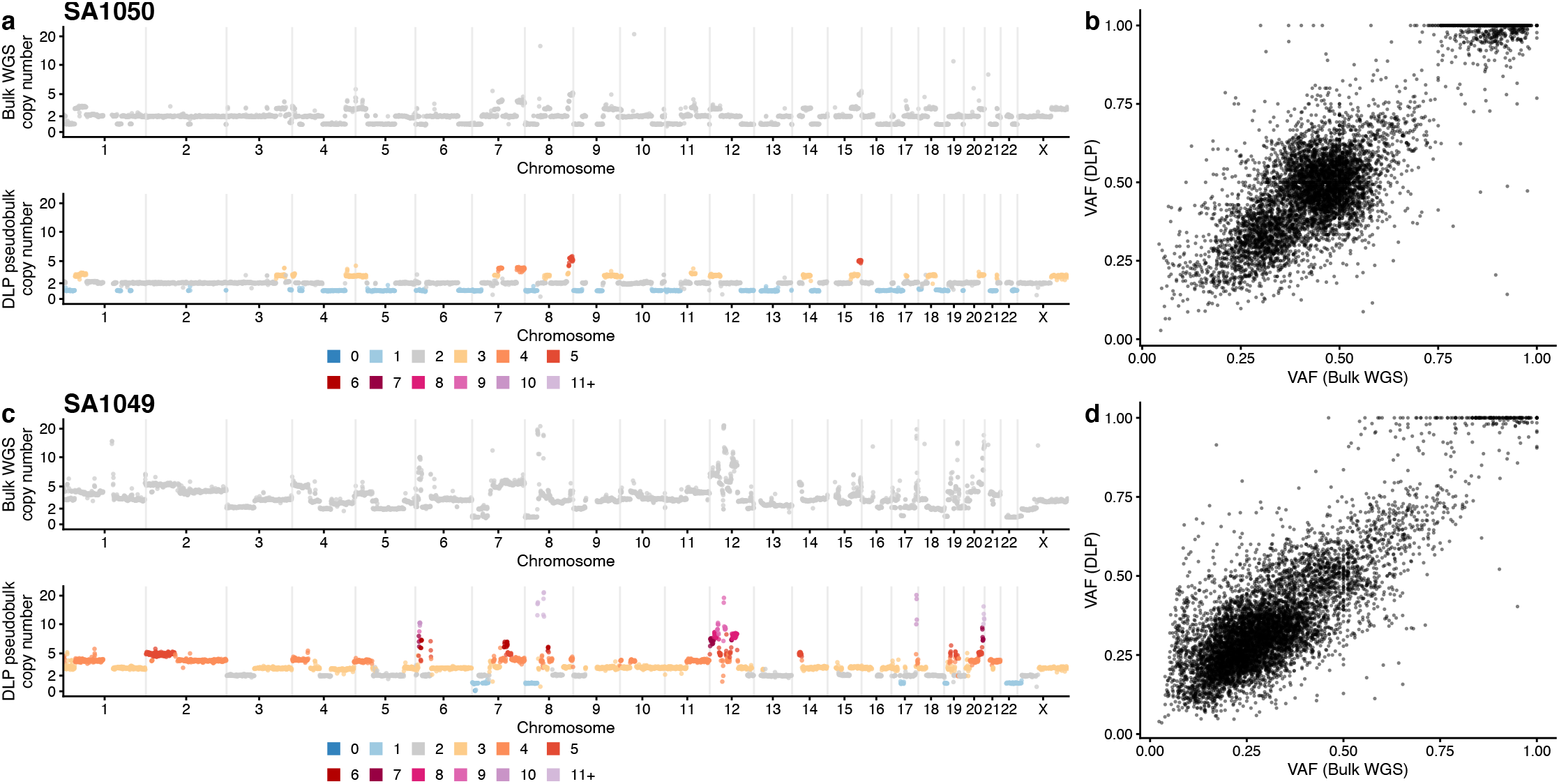
Comparison of copy number profiles and SNVs inferred from bulk WGS vs pseudobulk single cell DLP. a) Top: copy number in 500kb inferred from bulk WGS using Remixt, bottom: average copy number profile for all cells in SA1050. b) Variant allele fraction in bulk WGS vs DLP. c) and d) are equivalent to a) and b) but using sample SA1049.

**Extended Data Fig. 10.**
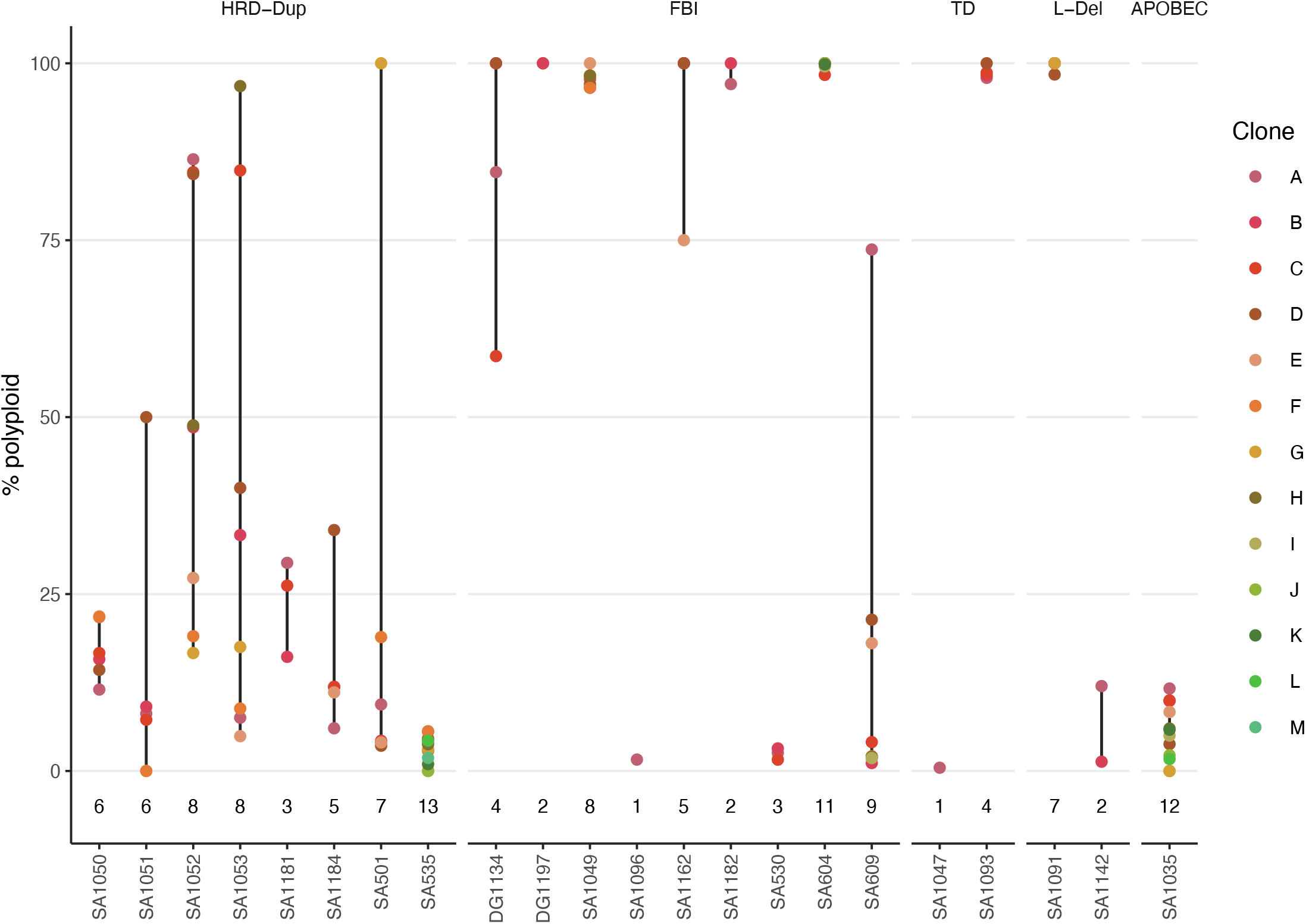
Cell polyploidy percentages for each clone in each dataset, split by signature type. # clones indicated below the data points. Minimum and maximum extents are indicated with black vertical lines. Clone colours are shared between datasets, but do not indicate any relationship between clones of different datasets.

**Extended Data Fig. 11.**
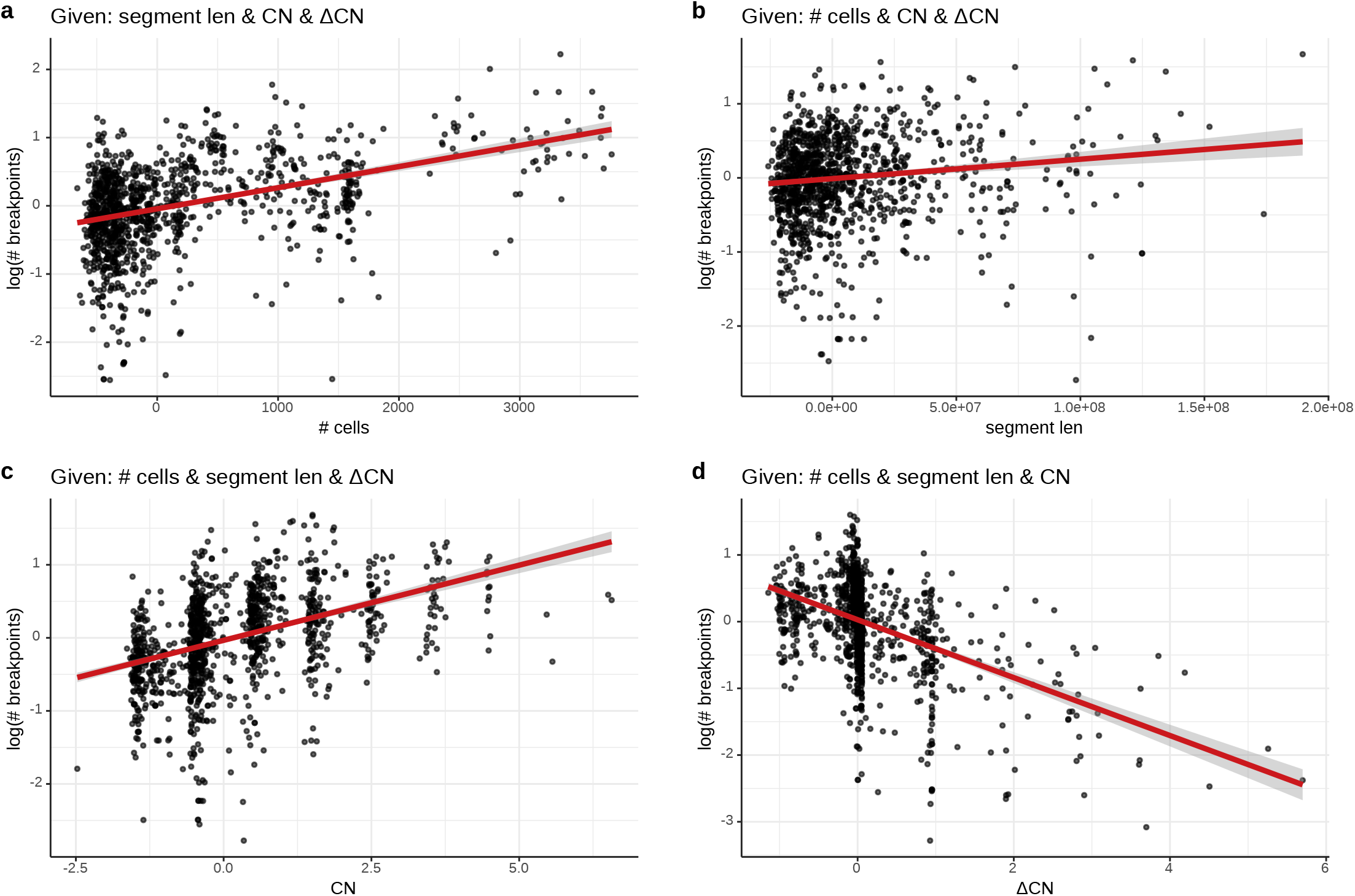
Partial regression plots showing relationship between the # cell breakpoints for an event and a covariate given the other covariates in a robust linear model. The conditioning variables are indicated in the panel titles. Robust linear model fits and standard error indicated with red lines and gray bands.

## Supplemental Tables

**Supplementary Table 1** HGSC and TNBC bulk sequencing library mutation signatures.

**Supplementary Table 2** Sample signature probabilities for HGSC and TNBC bulk sequencing libraries.

**Supplementary Table 3** Bulk sequencing sample clusters from mutation signature probabilities.

**Supplementary Table 4** List of induced mutations and CRISPR guides used in 184-hTERT cells. *TP53, BRCA1* or *BRCA2* mutations in tab 1 were induced by transduction with a CRISPR/Cas9 nuclease and one of the guides listed in tab 2. Mutations were determined by Sanger sequencing, recovered in DLP+ and protein knockout confirmed by western blotting.

**Supplementary Table 5** Per-cell level DLP+ sequencing statistics for 184-hTERT cell lines, HGSC, and TNBC tumours.

**Supplementary Table 6** Dataset level DLP+ sequencing statistics for 184-hTERT cell lines, HGSC, and TNBC tumours.

**Supplementary Table 7** Ploidy statistics for each cell included in the dataset.

**Supplementary Table 8** Chromosomal missegregation events for each cell included in the dataset.

**Supplementary Table 9** Copy number segments within chromosomes relative to cell ploidy.

**Supplementary Table 10** Gain and loss segments for dataset consensus copy number profiles relative to cell ploidy.

**Supplementary Table 11** Gain and loss segments for cell copy number profiles relative to the consensus profile of the clone to which the cell belongs.

**Supplementary Table 12** Copy number variances for 500 kb HLAMP bins.

**Supplementary Table 13** Max/min clone cell copy ratios for HLAMP genes.

**Supplementary Table 14** Breakpoint serration scores for chromosomal regions for each tumour in the dataset.

**Supplemental Methods** Supplementary method descriptions for data not included in the main text.

## Acknowledgments

This project was generously supported by the BC Cancer Foundation, Cycle for Survival supporting Memorial Sloan Kettering Cancer Center, the Terry Fox Research Institute, the CRUK Grand Challenge program, the Center for Excellence in Genome Sciences NIH grant (NIH 1RM1 HG01011014), Susan G Komen Breast Cancer Foundation leadership grant (GC260914) and the MSK SPORE in Genomic Instability in Breast Cancer Core grant (P50 CA247749-01) and the MSK Cancer Center Support Grant/Core Grant (P30 CA008748). JSR-F and BW are funded in part by Breast Cancer Research Foundation and National Institutes of Health/ National Cancer Institute P50 CA247749 01 grants. BW is funded in part by a Cycle for Survival grant. A.C.W. was supported by an NIH T32 training grant given to the Tri-Institutional Computational Biology and Medicine PhD Program. YF was supported by a BC Cancer Studentship.

## Author information

These authors contributed equally: Tyler Funnell and Ciara O’Flanagan.

## Contributions

SPS and SA: project conception and oversight, manuscript writing, senior responsible authors; TF, COF, MW, NR: manuscript writing and editing; COF, PE, FK, HL, TA, SL, AZ, JL, TM, JT, YF, VC: tissue procurement, biological substrates, knockout cell line generation and validation, data generation. COF, JB, BW, JB: single cell sequencing; TF, MW, JD, AM, DA, AW, NC, FU, DZ, ADCP: computational biology, data analysis; DL, SB, JP, DG, AM, SL, EH, VB: data processing, visualization; BW, SK, JR: pathology; SM: statistical, survival analysis; DY, HX, YL: discussion; RM: genome sequencing. All authors read and approved the final manuscript.

## Ethics declarations

BW reports ad hoc membership of the advisory board of Repare Therapeutics, outside the scope of this study. JSR-F reports receiving personal/consultancy fees from Goldman Sachs, REPARE Therapeutics, Paige.AI and Eli Lilly, membership of the scientific advisory boards of VolitionRx, REPARE Therapeutics and Paige.AI, membership of the Board of Directors of Grupo Oncoclinicas, and ad hoc membership of the scientific advisory boards of Roche Tissue Diagnostics, Ventana Medical Systems, Novartis, Genentech and InVicro, outside the scope of this study. SPS and SA are shareholders and consultants of Canexia Health.

