## Supplementary Methods for "The impact of mutational processes on structural genomic plasticity in cancer cells"

### Human mammary epithelial cell line generation and culture

The WT human mammary epithelial cell line 184-hTERT L9 (SA039) and isogenic 184-hTERT *TP53* KO (SA906) cell line, generated from 184-hTERT L9, were cultured as previously described [1–3] in MEBM (Lonza) supplemented with the SingleQuots kit (Lonza), 5 µg/ml transferrin (Sigma-Aldrich) and 10uM isoproterenol (Sigma-Aldrich). Additional truncation mutations (**Supplementary Table 4**) of BRCA1 (SA1054 : c.[427_441+36delGAAAATCCTTCCTTGGTAAAACCATTTGTTTTCTTC];[437_441+8delCC TTGGTAAAACC])

and BRCA2 (SA1056 : c.[6997delG];[6997_6998delGT]; SA1188 : c.[6997_6999delGT];[=]; SA1055 : c.[3507_3522delinsGA];[3509_3520delinT]) (hg19) were introduced by CRISPR/Cas9 nuclease (pX330 hSpCas9) with an RFP reporter gene using Mirus TransIT LT1 transfection (Mirus Bio). Clonal populations were generated by flow sorting and propagating single RFP positive cells. Mutations were verified by TOPO cloning and Sanger sequencing of both alleles for genotypes, protein expression by western blotting and absence of off-target effects by sequencing of the top hits. SNV positions from Sanger sequencing data were annotated with information from GENCODE version 19[4]. Variant sequence and position was used to annotate variant calls with records from Clinvar 20200206_data_release[5] and COSMIC version 91[6]. Although multiple BRCA2 homozygous loss of function alleles could be derived from 184-hTERT$^{p53-/-;BRCA2+/-}$ intermediates, only a single homozygous BRCA1 allele was retrieved from 184-hTERT$^{p53-/-;BRCA1+/-}$ 119 clones screened, emphasizing that even with a p53 deletion, full loss of BRCA1 is initially negatively selected.

### Human patient sample acquisition and consent

Samples were acquired with informed consent, according to procedures approved by the Ethics Committees at the University of British Columbia. Breast cancer patients undergoing diagnostic biopsy or surgery were recruited and samples collected under protocols H06-00289 (BCCA-TTR-BREAST), H11-01887 (Neoadjuvant Xenograft Study), H18-01113 (Large scale genomic analysis of human tumours) or H20-00170 (Linking clonal genomes to tumour evolution and therapeutics). HGSC samples were obtained from women undergoing debulking surgery under protocols H18-01652. Banked HGSC and TNBC specimens were obtained at Memorial Sloan Kettering Cancer Center following Institutional Review Board (IRB) approval and patient informed consent (protocols 15-200 (HGSC) and 18-376

(TNBC)). HGSC and TNBC clinical assignments were according to American Society of Clinical Oncology guidelines for ER, PR and HER2 positivity.

**Xenografting**

Patient tumour fragments were chopped finely with scalpels and mechanically disaggregated for one minute using a Stomacher 80 Biomaster (Seward Limited, Worthing UK) in 1 ml cold DMEM/F-12 with Glucose, L-Glutamine and HEPES (Lonza 12-719F). 200μl of medium containing cells/organoids from the resulting suspension was used equally for transplantation in 4 mice. The remaining tissue fragments were cryopreserved viably in DMEM/F12 supplemented with 47% FBS and 6% DMSO. Tumours were transplanted in mice as previously described[7] in accordance with SOP BCCRC 009. Female NOD/SCID/IL2rγ −/− (NSG) and NOD/Rag1−/−Il2rγ −/− (NRG) mice were bred and housed at the Animal Resource Centre (ARC) at the British Columbia (BC) Cancer Research Centre. For subcutaneous transplants, mechanically disaggregated cells and clumps of cells were resuspended in 150 - 200 μl of a 1:1 v/v mixture of cold DMEM/F12: Matrigel (BD Biosciences, San Jose, CA, USA). 8-12 week old mice were anesthetized with isofluorane and the mechanically disaggregated cell/clump suspension was transplanted under the skin on the left flank using a 1 ml syringe and 21 gauge needle. The animal care committee and animal welfare and ethical review committee, the University of British Columbia (UBC), approved all experimental procedures.

**Tissue processing**

Xenograft-bearing mice were euthanized when the size of the tumours approached 1000 mm3 in volume. The tumour material was excised aseptically and processed as described for primary tumour. Briefly, tumours were finely chopped, gently paddle blended and released single cells and fragments were viably frozen in DMEM supplemented with 47% FBS and 6% DMSO.

**Whole genome sequencing**

Genomic DNA was extracted from frozen tissue fragments using the DNeasy Blood and Tissue kit (Qiagen) and constructed libraries for whole genomes of 309 tumour-normal pairs were sequenced on the Illumina HiSeq X, according to Illumina protocols, generating 100bp paired-end reads for an estimated coverage of sequencing between 40X (normal) and 80X (tumour). Sequenced reads were aligned to the human reference GRCh37 (hg19) using BWA-MEM.

**Generation of single cell suspensions and nuclei for single cell DNA sequencing**

Viably frozen aliquots of patient tissues and PDX tumours were thawed and either homogenized and lysed using Nuclei EZ Buffer (Sigma) or enzymatically dissociated using a collagenase/hyaluronidase 1:10 (10X) enzyme mix (STEM CELL technologies), as described previously[3,8]. Cells and nuclei were stained with CellTrace CFSE (Life Technologies) and LIVE/DEAD Fixable Red Dead Cell Stain (ThermoFisher) in a 0.04% BSA/PBS (Miltenyi Biotec 130-091-376) incubated at 37C for 20 minutes. Cells were pelleted and resuspended in 0.04% BSA/PBS. This single cell suspension was loaded into a contactless piezoelectric dispenser (Cellenone or sciFLEXARRAYER S3, Scienion) and spotted into the open nanowell arrays (SmartChip, TakaraBio) preprinted with unique dual index sequencing primer pairs. Occupancy and cell state were confirmed by fluorescent imaging and wells were selected for single cell copy number profiling using the DLP+ method (Laks et al, 2019). Briefly, cell dispensing was followed by enzymatic and heat lysis. After cell lysis, tagmentation mix (14.335 nL TD Buffer, 3.5 nL TDE1, and 0.165 nL 10% Tween-20) in PCR water were dispensed into each well followed by incubation and neutralization. Final recovery and purification of single cell libraries was done after 8 cycles of PCR. Pooled single-cell libraries were analyzed using the Aglient Bioanalyzer 2100 HS kit. Libraries were sequenced at UBC Biomedical Research Centre (BRC) in Vancouver, British Columbia on the Illumina NextSeq 550 (midor high-output, paired-end 150-bp reads), or at the Genome Sciences Centre on Illumina HiSeq2500 (paired-end 125-bp reads) and Illumina HiSeqX (paired-end 150-bp reads). The data was then processed through a quantification and statistical analysis pipeline (Laks et al, 2019).

**Bulk whole genome data processing**

SNV and SV calls for 121 HGSC samples were acquired from Wang *et al.*[9]. For new samples, reads were aligned to the hg19 reference genomes using BWA-MEM. Processing proceeded as per Wang *et al.*[9] to maintain consistency.

SNVs were called with MutationSeq[10] (probability threshold = 0.9) and Strelka[11]. The intersection of calls from these methods were retained, however SNVs falling in blacklist regions were removed. The blacklist regions include the UCSC Genome Browser Duke and DAC blacklists, and those in the CRG Alignability 36mer track that had >2 mismatched nucleotides. SNVs were then annotated with SnpEff[12] for variant impact. SNV positions were annotated as for 184-hTERT lines.

SVs were called using deStruct[13] and LUMPY[14], and breakpoints called by both methods were retained. We then filtered events with the following criteria: any breakpoints fell in the blacklists described above, ≤30 bp inter-breakpoint distance, <1000 bp deletion, any breakpoints with <5 supporting reads in the tumour sample or any read support in the matched normal sample.

**HGSC & TNBC metacohort signature analysis**

Signatures analysis performed according to Funnell *et al.*[15] The MMCTM model was run on the sample SNV and SV count matrices. The number of signatures to estimate in the HGSC and TNBC integrated cohort was chosen by running the above fitting procedure for *k*=2–16 for both SNV and SV signatures with the number of restarts set to 500, where *k* is the number of signatures. We performed this step on approximately half the mutations in each sample, then computed the average per-mutation log-likelihood on the other held-out half of the mutations. The elbow curve method on log-likelihood values was used to select the final number of signatures to fit to the entire dataset.

To estimate MMCTM parameters on the full dataset, α hyper-parameters were set to 0.1. The model was initially fit to the data 1500 times. Each restart was run for a maximum of 1000 iterations or until the relative difference in predictive log likelihood on the training data was $<10^{-4}$ between iterations. The restarts with the best predictive log likelihoods for SNVs and SVs were selected as seeds for the final fitting step. The model was again fit to the data 1500 times. The model parameters for each restart were set to the parameters of the optimal models from the previous step described above, then run for a maximum of 1000 iterations or until the relative difference in predictive log likelihood on the training data was $<10^{-5}$ between iterations. The restart with the best mean rank of the SNV and SV predictive log likelihoods from this round was selected as the final model.

MMCTM estimated SNV signatures were matched to COSMIC signatures by solving the linear sum assignment problem for cosine distances between the MMCTM and COSMIC signatures using the clue R package[16]. An SNV signature matched to SBS12 was manually reassigned to SBS26 (MMRD) as there was also high cosine similarity to SBS26 (0.893, *vs.* 0.897 for SBS12), activity of this signature was enriched in a cluster of patients who were also enriched for another MMRD associated SNV signature, while SBS12 is mainly found in liver and biliary cancers[17].

**HGSC and TNBC patient survival analysis**

For each patient, the number of days between diagnosis and death, recurrence, or last follow-up were collected. For both HGSC and TNBC, patients were segregated into groups, and a Kaplan-Meier curve was fit for each group. Each cancer type was analyzed separately and in two distinct grouping schemes. First, patients were split into HRD and "Other" groups, where the HRD group included patients whose cancers were identified as being in either the HRD-Dup or HRD-Del groups, and the "Other" group included all other patients. Next, patients were grouped into these signature types: HRD-Dup, HRD-Del, FBI, TD, L-Del, APOBEC. This assignment is based on the following mapping of patient clusters to signature type: HRD-Dup (1), HRD-Del (3), FBI (2,7,8,9), TD (4), L-Del (11), APOBEC (5,6).

**DLP+ whole genome sequencing quantification and analysis**

Single cell copy number, SNV and SV calls were generated using the approach outlined in Laks *et al.*[8], except that BWA-MEM[18] was used to align DLP+ reads to the hg19 reference genome. 184-hTERT single cell sequencing targeting approximately 2 million reads per cell produced 4337 high quality single cell genomes in total (**Extended Data Fig. 8**) with median 0.04X coverage (IQR range: 0.03), and cancer single cell sequencing produced median 1.69 million reads per genome (median 0.05X coverage, IQR 0.04). The genome was segregated into 500 kb bins, and GC-corrected read counts were calculated for each bin. These read counts were then input into HMMCopy[19] to produce integer copy number states for each bin.

To detect SNVs and SVs in each dataset, reads from all cells in a DLP+ library were merged to form "pseudobulk" libraries. SNV calling was performed on these libraries individually using MutationSeq[10] (probability threshold = 0.9) and Strelka[11]. Only SNVs detected by both methods were retained. For each dataset, the union of SNVs was aggregated, then for each cell and each SNV, the sequencing reads of that cell were searched for evidence of that SNV. SV calling was performed in a similar manner, by forming pseudobulk libraries, then running LUMPY[11,14] on each pseudobulk library.

**DLP+ data filtering**

Cells were retained for further analysis if the cell quality was at least 0.75[8], and they passed both the s-phase and contamination filters. The contamination filter uses FastQ Screen[20] to tag reads as matching human, mouse, or salmon genomes. If >5% of reads in a cell are tagged as matching the mouse or salmon genomes, then the cell is flagged as contaminated. The s-phase filter uses the cell cycle state Random Forest classifier from Laks *et al.*[8] and removes cells where s-phase is the most probable state. The HGSC and TNBC cells were also filtered to remove small numbers of contaminating diploid cells.
A final cell filtering step was performed to remove putative early and late S-phase cells that passed the initial S-phase filter. This involved building a cell phylogeny with Sitka[21] and manually identifying the minimal phylogeny branches in which the cycling cells have been clustered. The cells in these branches were then removed.

We removed potentially problematic genome bins from our copy number results that had a mappability score of 0.99 or below, or were contained in the ENCODE hg19 blacklist[22].
SNV and SV calls from pseudobulk libraries were further post-processed according to Wang *et al*[9]**.**

**Identifying clones in DLP+ WGS by clustering copy number profiles**

For most datasets, clones were detected by first using UMAP on per-cell GC-corrected read count profiles, producing a 2-dimensional embedding of the cell profiles. We then ran HDBSCAN on the 2-dimensional embedding from UMAP to detect clusters of cells with similar copy number profiles.
UMAP was generally run with the following settings: n_neighbors = 100, min_dist = 0.0, and metric = "correlation". HDBSCAN was generally run with settings: min_samples = 5, min_cluster_size = 20, approx_min_span_tree = False, gen_min_span_tree = True, and allow_single_cluster = True. Dataset specific exceptions are as follows:

| | n_neighbors | min_samples | min_cluster_size |
|---|---|---|---|
| DG1134 | 40 | | 10 |
| SA1047 | 40 | 1 | |
| SA1091 | 3 | 10 | 15 |
| SA1093 | 25 | | |
| SA1096 | 10 | 1 | |
| SA1162 | 25 | 1 | 10 |
| SA1181 | 5 | | |
| SA1182 | 10 | | |
| SA1184 | 25 | | |
| SA535 | 12 | | |
| SA609 | 12 | 1 | 30 |
| SA1054 | 5 | | |

If HDBSCAN only finds a single cluster then the procedure stops. Otherwise HDBSCAN is run again with the additional setting of cluster_selection_epsilon = 0.2.

**Calculating cell ploidy**

Cell ploidy is calculated by taking the most common copy number state. Copy number states are those determined by HMMCopy.

**Identifying missegregated chromosomes**

The approach taken to identify putative chromosome missegregation events is similar to Laks *et al*[8]. Cells were split into groups corresponding to their clones. Clone copy number profiles were generated for each clone. Cells with ploidy not equal to the clone consensus profile were normalized to match the clone ploidy. Cell copy number profiles were compared to the clone copy

number profile for the matching clone to which the cell belongs. The result was assignment of an offset value for each genomic bin in each cell, that represented the copy number difference between cell and clone-level consensus profile. For each chromosome in each cell, if a particular copy number difference (*i.e.* -1, 1, etc.) represented at least 75% of the chromosome then we labelled that chromosome as having a missegregation event.

**Identifying CNA segments**

Gain and loss segments in each cell were found by comparing the copy number state in each 500 kb bin to that cell's ploidy. Copy number above ploidy was labelled as a gain, likewise copy number below ploidy was labelled as a loss. Gain and loss segments are a set of consecutive bins with the same gain/loss label. Segments ≤$1.5e^6$ kb were excluded to reduce segments potentially resulting from noise in the HMMCopy copy number states.

**Computing serriform variability scores in CNA breakpoints**

For each dataset, consensus copy number profiles were generated for each clone. Copy number segments were identified as above for each consensus profile. Copy number segments were then identified for single cell copy number profiles. The copy number profiles of each cell were normalized so that the adjusted cell ploidy matches the ploidy of the clone to which the cell belongs using the following formula:

$$cell\_state = cell\_state/cell\_ploidy * clone\_ploidy$$

Cell copy number segments were matched to segments in the clone copy number profile as follows: for each segment in the clone copy number profile, inspect the copy number states of the adjacent segments. If the segment state was less than both adjacent states, then only cell segments whose state was less than both of the two adjacent clone segment states could be matched to that segment. If the clone segment state was higher than both adjacent states, only cell segments whose state was higher than both of the adjacent clone segment states could be matched to that segment. If the clone segment state was in between the two adjacent states, only cell segments whose state was in between the two adjacent clone segment states could be matched to that segment. Finally, each cell segment was matched to the compatible clone segment that it overlapped the most, where compatibility means that the cell segment state met the criteria described above and the cell belonged to the relevant clone.

Next, clone segment breakpoints were aggregated across all clones. For each breakpoint, matched cell copy number segments were then queried for the number of unique cell segment breakpoint coordinates. This value was then log-transformed and corrected for the number of cells with that breakpoint, the length and copy number of the adjacent clone copy number profile segment with the highest copy number, and the difference in copy number between those adjacent states. The correction was performed by fitting a robust linear model with the log-transformed number of unique cell breakpoints as the dependent variable and the other values as independent variables, then taking the residuals and dividing by their standard deviation as the final serration score. Breakpoints whose higher-copy adjacent segments were longer than 10 Mbp were retained, while those with fewer than 100 cells were removed to retain only those breakpoints for which serration could be reliably computed. As a result, the DG1134 and DG1197 datasets (both FBI cases, see **Fig. 4**) were not included in comparisons as they have fewer than 100 cells each.

Highly serriform breakpoints were found by fitting a Normal distribution to all scores and choosing a cut off of the distribution mean + 1.5 s.d. Highly serriform breakpoints had scores above this threshold.

Clone-level serration was computed using a similar procedure as described above, except that cell breakpoint locations were first separated into groups according to cell clone assignments, rather than grouping all cell breakpoints across all cells in a dataset.

For each event, divergent breakpoint distances were determined by first identifying “mode” positions shared by >5% of cells, then calculating the distance between each cell breakpoint position and the nearest mode position. Distances >1 were classified as divergent and Beta distribution parameters were fit to these values. An expected null distribution of distances was generated by randomizing the locations of the divergent breakpoints within the event region and calculating distances to mode positions as before.

**Comparison of high-level amplification (HLAMP) copy number variance**

High level amplifications were identified by first selecting 500 kb genomic bins where at least 10 cells have raw copy number (adjusted per-bin read counts) of at least 10. Copy number variance for each bin was calculated using the raw copy number that was adjusted for cell ploidy and cell clone by first dividing the copy number by the cell ploidy, then subtracting the mean clone raw copy number. The cell ploidy is the most common hmmcopy copy number state as described above, and the mean clone copy number is computed for each bin in each clone across all cells in that clone. Mean HLAMP copy number variance was calculated for each dataset across all HLAMP bins, and these values were compared between signature type dataset groups.

**Immunoblotting**

184-hTERT cells were lysed directly in 1x Laemmli buffer supplemented with 7.5% beta mercaptoethanol and proteins were denatured at 95C for 15 mins. Protein from 250,000 cells was resolved on a 4-15% acrylamide gel (Biorad) or 3-8% Tris-acetate acrylamide gel (Novex) and transferred to a nitrocellulose membrane with Towbin transfer buffer overnight at 30V at 4C. Blots were blocked with 5% Milk in TBST for 1 hr and incubated overnight at 4C with mouse anti-p53 (Santa Cruz SC-126, 1:500 in 5% BSA), mouse anti-BRCA1 (Santa Cruz SC-6954, 1:200 in 5% BSA), mouse anti-BRCA2 (Millipore OP95, 1:200 in 5% BSA) or goat anti-GAPDH (SC-48166, 1:500 in 5% BSA). Blots were washed 5 times for 5 mins in TBS-T and incubated with anti-goat HRP-conjugated secondary antibody (Abcam ab6721, 1:5000 in 5% BSA) for 1 hr at room temperature, washed 5 times for 5 mins and signal imaged using Immobilon Western Chemiluminescent HRP Substrate (MilliporeSigma, cat#WBKL20500) and the ImageQuant LAS 4000 (GE Healthcare) using the ImageQuant TL software.

**184-hTert cell line mutation verification**

Genomic DNA was extracted from 184-hTERT cell lines and regions of interest of BRCA1 or BRCA2 were amplified by PCR. Amplicons were inserted into a pCR-TOPO vector and transformed into *E.Coli* using the TOPO TA cloning kit (Thermo Fisher). Colonies were selected, DNA purified by Purelink Quick Plasmid Miniprep kit (Thermo Fisher) and sequenced by Sanger sequencing to assess CRISPR-induced mutations. Mutations were detected in DLP+ sequencing data by visualizing in the integrative genomics viewer (IGV) tool.

**Drug sensitivity**

184-hTERT cells were seeded at a concentration of 500 cells per well into a 384-well plate (n=5 replicates) and incubated for 24hrs before being treated with increasing concentrations of cisplatin for 96hrs. Cells were lysed and viability measured using Cell Titre Glo (Promega) according to the manufacturer's instructions. IC50 values were determined by non-linear regression analysis of resulting curves.

**Cox proportional hazards modeling**

We selected 6 signatures (S-Dup, S-Del, FBI/Inv, L-Dup, APOBEC2, and L-Del) that represented the enriched signatures in the 6 major patient strata (*i.e.* those with at least 10 cases: HRD-Dup, HRD-Del, FBI, TD, APOBEC, L-Del). The standardized activities of these signatures and the age at diagnosis for each patient were input into a Cox proportional hazards model with either overall or progression free survival as the outcome. For TNBC cases, tumour grade, size, and node status were included as additional covariates.

**Statistical Tests**

Statistical tests used were two-tailed unequal-variance t-tests unless otherwise specified: log-rank tests were used for comparing survival curves; Wilcoxon rank sum two-tailed tests were used for comparing segment lengths, segment counts, copy variances, bin counts, and breakpoint counts; Kolmogorov-Smirnov two-tailed tests were used to compare continuous distributions. ANOVA was used to compute p-values for comparisons of clone breakpoint serration. P-values from multiple comparisons were corrected using the Benjamini & Hochberg method.

**Code availability**

MMCTM method: https://github.com/shahcompbio/MultiModalMuSig.jl
DLP+ single cell WGS pipeline: https://github.com/shahcompbio/single_cell_pipeline
Bulk WGS pipeline: https://github.com/shahcompbio/wgs


1. Burleigh, A. *et al.* A co-culture genome-wide RNAi screen with mammary epithelial cells reveals transmembrane signals required for growth and differentiation. *Breast Cancer Res.* **17**, 4 (2015).
2. Horikawa, I. *et al.* Downstream E-Box--mediated Regulation of the Human Telomerase Reverse Transcriptase (hTERT) Gene Transcription: Evidence for an Endogenous

Mechanism of Transcriptional Repression. *Mol. Biol. Cell* **13**, 2585–2597 (2002).

3. Salehi, S. *et al.* Single cell fitness landscapes induced by genetic and pharmacologic perturbations in cancer. *Cold Spring Harbor Laboratory* 2020.05.08.081349 (2020) doi:10.1101/2020.05.08.081349.
4. Frankish, A. *et al.* GENCODE reference annotation for the human and mouse genomes. *Nucleic Acids Res.* **47**, D766–D773 (2019).
5. Landrum, M. J. *et al.* ClinVar: improving access to variant interpretations and supporting evidence. *Nucleic Acids Res.* **46**, D1062–D1067 (2018).
6. Tate, J. G. *et al.* COSMIC: the Catalogue Of Somatic Mutations In Cancer. *Nucleic Acids Res.* **47**, D941–D947 (2019).
7. Eirew, P. *et al.* Dynamics of genomic clones in breast cancer patient xenografts at single-cell resolution. *Nature* **518**, 422–426 (2015).
8. Laks, E. *et al.* Clonal Decomposition and DNA Replication States Defined by Scaled Single-Cell Genome Sequencing. *Cell* **179**, 1207–1221.e22 (2019).
9. Wang, Y. K. *et al.* Genomic consequences of aberrant DNA repair mechanisms stratify ovarian cancer histotypes. *Nat. Genet.* **49**, 856–865 (2017).
10. Ding, J. *et al.* Feature-based classifiers for somatic mutation detection in tumour-normal paired sequencing data. *Bioinformatics* **28**, 167–175 (2012).
11. Saunders, C. T. *et al.* Strelka: accurate somatic small-variant calling from sequenced tumor-normal sample pairs. *Bioinformatics* **28**, 1811–1817 (2012).
12. Cingolani, P. *et al.* A program for annotating and predicting the effects of single nucleotide polymorphisms, SnpEff. *Fly* vol. 6 80–92 (2012).
13. McPherson, A., Shah, S. P. & Sahinalp, S. C. deStruct: Accurate Rearrangement Detection using Breakpoint Specific Realignment. *bioRxiv* 117523 (2017).
14. Layer, R. M., Chiang, C., Quinlan, A. R. & Hall, I. M. LUMPY: a probabilistic framework for structural variant discovery. *Genome Biol.* **15**, R84 (2014).
15. Funnell, T. *et al.* Integrated structural variation and point mutation signatures in cancer genomes using correlated topic models. *PLOS Computational Biology* vol. 15 e1006799

(2019).

16. Hornik, K. A CLUE for CLUster Ensembles. *J. Stat. Softw.* **14**, (2005).
17. Alexandrov, L. B. *et al.* The repertoire of mutational signatures in human cancer. *Nature* **578**, 94–101 (2020).
18. Li, H. Aligning sequence reads, clone sequences and assembly contigs with BWA-MEM. *arXiv [q-bio.GN]* (2013).
19. HMMcopy. http://bioconductor.org/packages/HMMcopy/.
20. Wingett, S. W. & Andrews, S. FastQ Screen: A tool for multi-genome mapping and quality control. *F1000Res.* **7**, 1338 (2018).
21. Dorri, F. *et al.* Efficient Bayesian inference of phylogenetic trees from large scale, low-depth genome-wide single-cell data. *Cold Spring Harbor Laboratory* 2020.05.06.058180 (2020) doi:10.1101/2020.05.06.058180.
22. Amemiya, H. M., Kundaje, A. & Boyle, A. P. The ENCODE Blacklist: Identification of Problematic Regions of the Genome. *Sci. Rep.* **9**, 1–5 (2019).
